# The ATP-exporting channel Pannexin-1 promotes CD8^+^ T cell effector and memory responses

**DOI:** 10.1101/2023.04.19.537580

**Authors:** Trupti Vardam-Kaur, Alma Banuelos, Maria Gabaldon-Parish, Bruna Gois Macedo, Caio Loureiro Salgado, Kelsey Marie Wanhainen, Maggie Hanqi Zhou, Sarah van Dijk, Igor Santiago-Carvalho, Angad S. Beniwal, Chloe L. Leff, Changwei Peng, Nhan L. Tran, Stephen C. Jameson, Henrique Borges da Silva

**Author notes:** Corresponding author and lead contact (H.BdS). These authors contributed equally to this manuscript (Trupti Vardam-Kaur, Alma Banuelos, Maria Gabaldon-Parish, Bruna Gois Macedo).

## Abstract

Sensing of extracellular ATP (eATP) controls CD8^+^ T cell function. Their accumulation can occur through export by specialized molecules, such as the release channel Pannexin-1 (Panx1). Whether Panx1 controls CD8^+^ T cell immune responses *in vivo*, however, has not been previously addressed. Here, we report that T cell-specific Panx1 is needed for CD8^+^ T cell responses to viral infections and cancer. We found that CD8-specific Panx1 promotes both effector and memory CD8^+^ T cell responses. Panx1 favors initial effector CD8^+^ T cell activation through extracellular ATP (eATP) export and subsequent P2RX4 activation, which helps promote full effector differentiation through extracellular lactate accumulation and its subsequent recycling. In contrast, Panx1 promotes memory CD8^+^ T cell survival primarily through ATP export and subsequent P2RX7 engagement, leading to improved mitochondrial metabolism. In summary, Panx1-mediated eATP export regulates effector and memory CD8^+^ T cells through distinct purinergic receptors and different metabolic and signaling pathways.

## Introduction

Cytotoxic CD8^+^ T cells control viral infections and solid tumors ^1,2^. These cells, upon antigen priming, quickly activate intracellular signaling pathways that lead to effector function and clonal expansion ^2^. The expansion and differentiation of effector CD8+ T cells are controlled by a network of supporting cells and extracellular signals. Effector CD8^+^ T cells differentiate into either memory precursors (MP) cells or terminal effectors (TE), which perform distinct functions and have different fates during T cell contraction. While MP cells become long-lived memory CD8^+^ T cell subsets, TE cells display heightened effector function and mostly die during contraction ^2^, except for long-lived effector cells (LLECs) ^3,4^. Effector CD8^+^ T cells expansion and differentiation are controlled by a network of supporting cells, and extracellular signals ^2,5–7^. Various components of this network have been identified, including interactions with innate immune cells^8,9^ and cytokine sensing ^10–13^. Extracellular accumulation of metabolites, however, may also play an important role. Metabolites are often released as a byproduct of dead cells ^14^. However, they can sometimes be actively released via transmembrane channels ^15–17^.

One of the most well-described metabolite transporters is the homo-heptameric release channel Pannexin-1 (Panx1) ^18,19^. Through release of “find-me” signals such as the nucleotide ATP, Panx1 plays a critical role in the triggering of apoptotic cell clearance ^19–22^. Panx1 can also promote ATP export in healthy cells ^20,23–27^. Panx1 is expressed by many immune and non-immune cells, including T cells ^28,29^. Panx-1-mediated ATP export promotes the anti-allergic function of regulatory CD4^+^ T cells (Tregs) ^30^. Conventional CD4^+^ T cells or CD8^+^ T cells can also export ATP via Panx1, influencing their function ^28,31,32^. Panx1 also promote the export of many other metabolites smaller than 1 kDa ^19^ ^20^ ^21^. Metabolite export via Panx1, therefore, can regulate T cell function and its downstream physiological consequences. However, no comprehensive studies on the *in vivo* role of Panx1 in CD8^+^ T cells have been done.

Extracellular ATP (eATP) is sensed by immune cells via purinergic receptors, like the high-affinity ion channel P2RX4 or low-affinity ion channel P2RX7 ^33^. We and others have previously shown that P2RX7 is expressed by antigen-specific CD8^+^ T cells ^34–36^ and is crucial for the generation and survival of memory CD8^+^ T cells ^32,34,37^. In addition to Panx1 export, eATP can accumulate through infection-induced tissue damage ^38^. The role of P2RX7 for memory CD8^+^ T cell survival raises an interesting conundrum: these cells rely on consistent eATP signaling, yet the release of ATP from tissue damage is likely absent after the pathogen is gone. In this work, we investigated the role of Panx1 in the development of antigen-specific CD8^+^ T cell responses to viral infection. We found that Panx1 favors both the expansion of effector CD8^+^ T cells and the survival of memory CD8^+^ T cells. Our data suggests that Panx1 promotes effector CD8^+^ T cells through a two-step process: following TCR priming, Panx1-mediated eATP export promotes initial CD8^+^ T cell activation in a P2RX4-dependent way. At a later stage, eATP is insufficient to restore activation; Panx1-mediated activation leads to the accumulation of extracellular lactate which is subsequently recycled into saturated phosphatidylinositol biosynthesis. In contrast, memory CD8^+^ T cells require Panx1 for survival through eATP export and subsequent sensing of it via P2RX7. The overarching role of Panx1 for CD8^+^ T cells is evidenced by its importance in promoting optimal control of melanoma tumors by CD8^+^ T cells, as well as the induction of CD8^+^ T cell-mediated graft-versus-host disease.

## Results

### T cell-specific Panx1 promotes effector CD8^+^ T cell accumulation

We first sought to define how Panx1 affects CD8^+^ T cell effector responses (gating strategy in **Figure S1**). Panx1 mRNA levels are expressed by antigen-specific CD8^+^ T cells (**Figure 1a, Figure S2a**). These channels are functional in CD8^+^ T cells, as ATP export was significantly decreased – and intracellular ATP levels are concomitantly increased – when Panx1 is inhibited by Trovafloxacin, an antibiotic with known Panx1-blocking function^22^ (**Figure S2b**). Panx1 can promote nucleotide export in apoptotic cells upon triggering by Caspase 3^22^. However, we did not find increased To-Pro3 uptake in WT CD8^+^ T cells in comparison to CD8^+^ T cells deficient for Panx1 (Panx1-KO) (**Figure S2c**), suggesting that Panx1 primarily drives metabolite export from healthy CD8^+^ T cells. Importantly, T cell-specific knockout of Panx1 does not affect CD8^+^ T cell thymic development (**Figure S2d**) or steady-state peripheral accumulation (**Figure S2e**).

**Figure 1.**
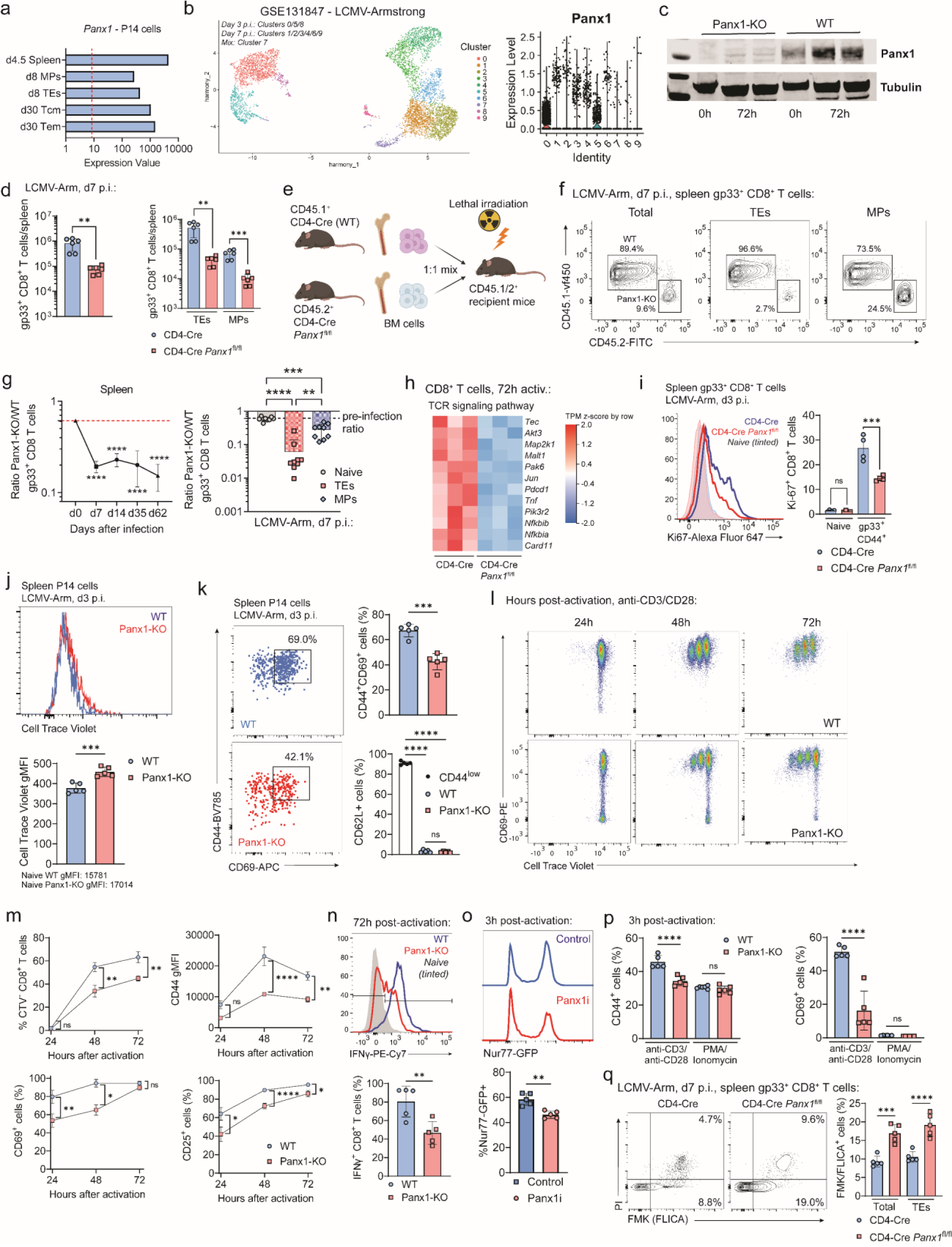
CD8^+^ T cell-specific Panx1 is required for T cell activation and effector differentiation. (**a**) *Panx1* mRNA expression from RNA-seq samples in the indicated subsets of LCMV-gp33 transgenic P14 CD8^+^ T cells. (**b**) Single-cell RNA-seq (scRNAseq) analysis of effector P14 cells showing UMAP cluster distribution (left) and expression of *Panx1* mRNA in each cluster (right). (**c**) Western Blot analysis of Panx1 expression in WT (CD4-Cre) or Panx1-KO (CD4-Cre *Panx1*^fl/fl^) CD8^+^ T cells at 0h or 72h after activation with anti-CD3/CD28 + IL-2. (**d**) WT (CD4-Cre) or CD4-Cre *Panx1*^fl/fl^ mice were infected with LCMV-Arm. Numbers of total (left) and TE, MP (right) spleen gp33^+^ CD8^+^ T cells at day 7 post-infection are shown. (**e-g**) A 1:1 mix of bone marrow cells from CD4-Cre (WT; CD45.1^+^) and CD4-Cre *Panx1*^fl/fl^ (Panx1-KO; CD45.2^+^) mice was transferred into lethally irradiated C57BL/6 mice (CD45.1/2^+^). After two months, BM chimeric mice were infected with LCMV-Arm. The red dotted lines show the Panx1-KO:WT ratios after two months of reconstitution. (**e**) Graphical scheme. (**f**) Representative flow cytometry plots showing WT and Panx1-KO total, TE and MP spleen gp33^+^ CD8^+^ T cells at day 7 post-infection. (**g**) Panx1-KO/WT ratios of total (left) and of naïve, TE and MP spleen gp33^+^ CD8^+^ T cells (right) at day 7 post-infection. (**h**) Heatmaps showing the expression of TCR signaling pathway genes from RNA-seq data of activated WT and Panx1-KO CD8^+^ T cells. (**i**) CD4-Cre or CD4-Cre *Panx1*^fl/fl^ mice were infected with LCMV-Arm and evaluated at 3 days after infection. Representative histograms (left) and average percentages (right) of Ki-67^+^ gp33^+^ CD8^+^ T cells are shown. (**j-k**) CD4-Cre (WT; CD45.1/2^+^) and CD4-Cre *Panx1*^fl/fl^ (Panx1-KO; CD45.2^+^) P14 cells were labeled with Cell Trace Violet and co-transferred into LCMV-infected recipient C57BL/6 (CD45.1^+^) mice. (**j**) Histograms showing Cell Trace Violet dilution (left) and graphs showing average gMFI values (right) are shown. (**k**) Representative flow cytometry plots showing expression of CD44 and CD69 are shown in the left. In the right, average percentages of CD44^+^CD69^+^ cells (above) and of CD62L^+^ cells (below) are shown. (**l-q**) WT (CD4-Cre) or Panx1-KO (CD4-Cre *Panx1*^fl/fl^) CD8^+^ T cells were activated *in vitro* (anti-CD3/CD28 + IL-2, and in some cases PMA/Ionomycin) for up to 72h. In some experiments (**o**), WT Nur77-GFP CD8^+^ T cells were activated in the presence of vehicle (PBS; Control) or Trovafloxacin (Panx1 inhibitor; Panx1i). (**l**) Representative flow cytometry plots showing expression of CD69 and dilution of Cell Trace Violet at 24h, 48h and 72h after activation. (**m**) Percentages of Cell Trace Violet (CTV) divided cells (top left), average gMFI values for CD44 (top right), and percentages of CD69^+^ (bottom left) and CD25^+^ (bottom right) cells at 24h, 48h and 72h after activation. (**n**) Representative histograms showing expression of IFNγ (top) and average percentages of IFNγ^+^ CD8^+^ T cells (bottom) at 72h after activation. (**o**) Representative histograms showing Nur77-GFP expression in control of Panx1i-treated cells (above), and average percentages of Nur77-GFP^+^ CD8^+^ T cells (below) at 3h after activation are shown. (**p**) Average percentages of CD44^+^ (left) and CD69^+^ (right) cells at 3h after activation with either anti-CD3/CD28 or PMA/Ionomycin are shown. (**q**) CD4-Cre and CD4-Cre *Panx1*^fl/fl^ mice were infected with LCMV-Arm; at day 7 after infection, the percentages of FMK/FLICA^+^ and PI^+^ cells were assessed in gp33^+^ CD8^+^ T cells. In the left, representative flow cytometry plots are shown; in the right, average percentages of FMK/FLICA+ cells are depicted. (**c**) Data representative of 2 independent experiments, n=4-5 per group. (**d-j, q**) Data from 2-3 independent experiments, n=5-12 per experimental group. (**k**) Each replicate is a pool of activated CD8^+^ T cells from three mice; n=3 replicates per experimental group. (**l-p**) Data from four independent experiments, n=7-11 per experimental group per time point. ns – not significant (p>0.05), *p<0.05, **p<0.01, ***p<0.001, ****p<0.0001, Unpaired t-test (**d, j-k, n-o**), One-way ANOVA with Tukey’s post-test (**d, g, i, k, p-q**), Two-way ANOVA with Bonferroni’s post-test (**g, m**).

Despite its presence in naïve CD8^+^ T cells, Panx1 mRNA and protein expression levels are increased in effector CD8^+^ T cells (**Figures 1b-c, S2f-g**). Given its inhibitory role, we used Trovafloxacin treatment of Lymphocytic Choriomeningitis Virus (LCMV)-infected mice (previously transferred with LCMV gp33-TCR transgenic CD8^+^ P14 cells) to test how Panx1 blockade affects the effector expansion of CD8^+^ T cells. At one-week post-infection, we observed significantly decreased numbers of splenic P14 cells (**Figure S3a**). Since multiple other immune and non-immune cells can express Panx1 (**Figure S3b**), we used T cell-specific KO systems to test the CD8^+^ T cell-specific role of Panx1. LCMV-infected T cell-specific Panx1-KO (CD4-Cre *Panx1*^fl/fl^) mice had significantly decreased numbers of spleen effector gp33-specific CD8^+^ T cells (**Figure 1d**, left). Both KLRG1^+^CD127^-^ terminal effectors (TE) and CD127^+^KLRG1^-^ memory precursors (MP) numbers were lower in CD4-Cre *Panx1*^fl/fl^ mice (**Figure 1d**, right). Similar results were found using alternative experimental systems: 1:1 WT:CD4-Cre *Panx1*^fl/fl^ mixed bone marrow chimeras infected with LCMV (**Figures 1e-g**), CRISPR-Cas9 deletion of Panx1 in P14 cells (sgPanx1; **Figures S4a-b**), and infection of WT versus CD4-Cre *Panx1*^fl/fl^ with influenza (**Figure S4c**) – despite no differences in the lung accumulation of influenza-specific CD8^+^ T cells being observed (**Figure S4d**).

We next measured which aspects of effector CD8^+^ T cell function rely on Panx1 expression. RNA-seq analysis of WT versus Panx1-KO CD8^+^ T cells activated *in vitro* for 72h showed decreased expression of many genes associated with the T cell receptor (TCR) signaling pathway in Panx1-KO CD8^+^ T cells (**Figure 1h**). Indeed, Panx1-KO CD8^+^ T cells displayed reduced proliferation *in vivo* (**Figures 1i-j**). In addition to proliferation defects, Panx1-KO CD8^+^ T cells from LCMV-infected mice expressed reduced levels of activation markers such as CD69 or CD44 (**Figure 1k**). *In vitro* proliferative capacity (**Figures 1l-m**), expression of CD69, CD44 or CD25 (**Figure 1m**) and production of IFN-γ (**Figure 1n**) by Panx1-KO CD8^+^ T cells was also impaired.

Panx1 acts as a release channel for metabolites such as eATP. Our data shows that Panx1-KO affects effector CD8^+^ T cell activation at a populational level, making it hard to distinguish between paracrine versus autocrine roles for Panx1. To discern between these two possibilities, we used 1:1 WT:Panx1-KO co-cultures of CD8^+^ T cells in *in vitro* activation assays (**Figure S4e**). As shown in **Figure S4f**, despite some paracrine rescue, the expression of CD69 and proliferation of Panx1-KO CD8^+^ T cells were still significantly lower than WT CD8^+^ T cells, even in 1:1 co-culture. These results, which agree with our 1:1 mixed bone marrow chimeras, suggest a dominant autocrine role for Panx1 for effector CD8^+^ T cells, despite some paracrine effects.

Panx1 is recruited to the immunological synapse after T cell priming ^28^. Therefore, Panx1 may promote the activation of CD8^+^ T cells in a TCR-dependent manner. Indeed, upregulation of TCR downstream signaling molecule Nur77 is significantly impaired in the presence of Trovafloxacin (Panx1i) (**Figure 1o**). Additionally, no defects in early CD8^+^ T cell activation were observed in Panx1-KO CD8^+^ T cells upon short-term stimulation with PMA and Ionomycin, which bypass the TCR signaling pathway (**Figure 1p**), further indicating a TCR-dependent role for Panx1. Finally, we measured if Panx1-KO would affect effector CD8^+^ T cell survival. As shown in **Figure 1q**, Panx1-KO effector CD8^+^ T cells had increased percentages of FLICA^+^ (PI^+^) apoptotic cells. Overall, our results show that Panx1 is necessary for the effector CD8^+^ T cell proliferation, activation, and resistance to apoptosis in response to acute viral infection.

### Panx1 regulates the intracellular metabolism of effector CD8^+^ T cells

Because the activation and effector differentiation of CD8^+^ T cells require sequential metabolic adaptations, we investigated how Panx1 controls the intracellular metabolism of CD8^+^ T cells. Naïve Panx1-KO CD8^+^ T cells showed no defects in mitochondrial respiration, mitochondrial mass, or membrane potential (**Figures 2a-b**). However, as early as 3h after stimulation and following early defects in activation, Panx1 deficiency negatively affected mitochondrial mass and membrane potential (**Figure 2c**, above), leading to sustained defects up to 72h post-activation (**Figure 2c**, below). After 72h of activation, Panx1-KO CD8^+^ T cells displayed reduced engagement of bioenergetic pathways needed for initial CD8^+^ T cell activation ^39^: aerobic glycolysis and mitochondrial respiration (**Figure 2d**). Similar defects in mitochondrial mass and membrane potential were observed in Panx1-KO CD8^+^ T cells at early time points *in vivo* (**Figure S5a**).

**Figure 2.**
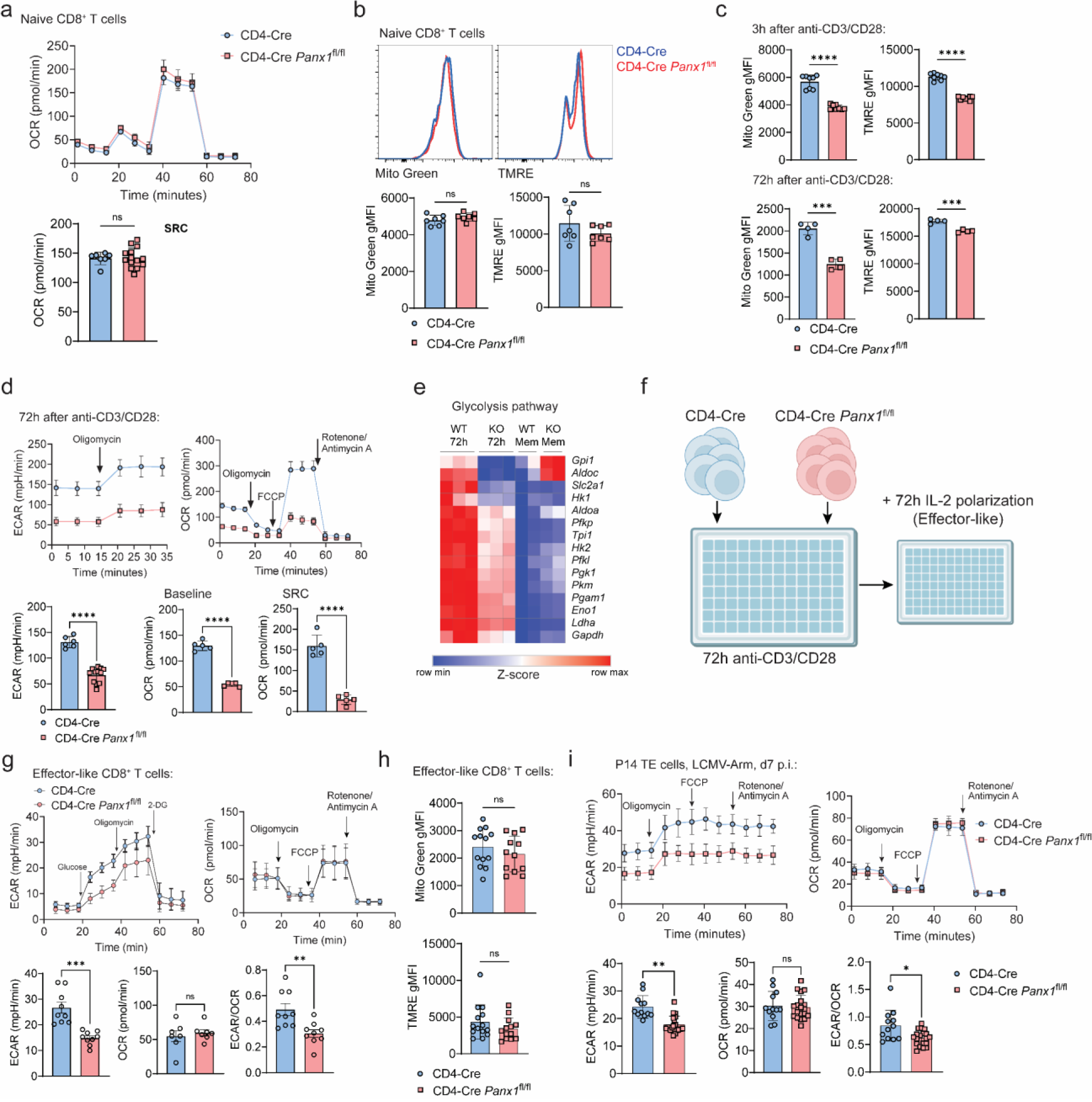
Panx1 controls the intracellular metabolism of effector CD8^+^ T cells. (**a-b**) Naïve CD4-Cre and CD4-Cre *Panx1*^fl/fl^ CD8^+^ T cells were analyzed for mitochondrial function. (**a**) Measurements of Oxygen Consumption Rate (OCR) levels after sequential addition of Oligomycin, FCCP and Rotenone/Antimycin A (top right); average OCR values at Spare Respiratory Capacity (bottom right, SRC; see methods) are shown. (**b**) Mitotracker Green (Mito Green) and TMRE; representative histograms are shown above, while average gMFI values are shown below. (**c-d**) CD4-Cre or CD4-Cre *Panx1*^fl/fl^ CD8^+^ T cells were activated *in vitro* (anti-CD3/CD28 + IL-2) for up to 72h. (**c**) Mito Green and TMRE average gMFI levels at 3h (above) and 72h (below) after activation are shown. (**d**) Measurements of Extracellular Acidification Rate (ECAR) levels after addition of Oligomycin (top left); measurements of Oxygen Consumption Rate (OCR) levels after sequential addition of Oligomycin, FCCP and Rotenone/Antimycin A (top right); average ECAR values at baseline (bottom left), average OCR values at baseline (bottom middle) and Spare Respiratory Capacity (bottom right, SRC; see methods) are shown. (**e**) Heatmap showing expression of genes from the Glycolysis pathway in 72h-activated CD8^+^ T cells; expression in memory-like CD8^+^ T cells (with additional IL-15 treatment – more on Figure 6) are shown in comparison. (**f-h**) CD4-Cre or CD4-Cre *Panx1*^fl/fl^ CD8^+^ T cells were activated *in vitro* (anti-CD3/CD28 + IL-2) for 72h, then further polarized into effector-like cells with IL-2 for additional 72h. (**f**) Experimental design. (**g**) Measurements of Extracellular Acidification Rate (ECAR) levels from effector-like CD8^+^ T cells after sequential addition of Glucose, Oligomycin and 2-DG (top left); in the top right, kinetics of OCR values. On the bottom, average baseline ECAR and OCR values, and the ratios between ECAR and OCR values from effector-like CD8^+^ T cells are shown respectively. (**h**) Average Mito Green and TMRE gMFI values for effector-like CD8^+^ T cells. (**i**) CD4-Cre and CD4-Cre *Panx1*^fl/fl^ P14 cells were transferred into C57BL/6 mice infected with LCMV-Arm. At day 7 post-infection, TE P14 cells were sorted. ECAR and OCR kinetics (top), ECAR values (bottom left), OCR values (bottom center) and ECAR/OCR ratios (bottom right) from sorted TE gp33^+^ CD8^+^ T cells are shown. (**a-d, f-i**) Data from 2-3 independent experiments, n=4-21 per experimental group. (**e**) Data from 2-3 biological replicates per experimental group (from n=5-8 mice per group). ns – not significant (p>0.05), *p<0.05, **p<0.01, ***p<0.001, ****p<0.0001, Unpaired t-test (**a-d, f-i**).

Following initial activation, effector CD8^+^ T cells must shift their metabolism towards preferential usage of aerobic glycolysis for terminal effector differentiation^40^. *In vitro*-activated CD8^+^ T cells for 72h displayed profound defects in mRNA expression of genes associated with aerobic glycolysis (**Figure 2e**). To understand how Panx1 regulates the metabolism of CD8^+^ T cells at this late activation stage, we used prolonged exposure of *in vitro*-activated CD8^+^ T cells to IL-2, mimicking effector-like conditions^41^ (**Figure 2f**). Effector-like CD8^+^ T cells exhibited reduced engagement of aerobic glycolysis but showed no defects in mitochondrial respiration, mass, or membrane potential (**Figures 2g-h**). To determine if Panx1 specifically affects the aerobic glycolysis of late effector CD8^+^ T cells, we sorted TE P14 cells from LCMV-infected mice (WT versus T cell-Panx1-KO) for extracellular flux analyses. Panx1-KO TE cells showed reduced glycolysis but normal mitochondrial respiration (**Figure 2i**). These results indicate that Panx1 plays a dominant role in promoting the metabolic adaptations required for initial CD8^+^ T cell activation (energetic metabolism increase) and effector CD8^+^ T cell full differentiation (induction of aerobic glycolysis).

### Panx1-KO defects in effector CD8^+^ T cells can be rescued by exogenous eATP and sodium lactate

Both 72h-activated (**Figure S2b**) and IL-2-induced effector-like CD8^+^ T cells (**Figure S5b**) export ATP via Panx1. Due to this well-established role of Panx1 and previous evidence *in vitro*^31,42^, we hypothesized that exogenous eATP could rescue the activation defects of Panx1-KO CD8^+^ T cells. Early (3h post-stimulation) activation defects are significantly restored: Nur77, CD69, CD25 and CD44 upregulation in Panx1-KO CD8^+^ T cells are all significantly restored by eATP addition to cultures (**Figures 3a-b, S5c**). Consistent with a previous study^42^, the eATP rescue of Panx1-KO activation is abolished in the presence of 5-BDBD, a P2RX4 inhibitor (**Figure S5c**). Indeed, P2RX4 deletion in CD8^+^ T cells phenocopied the defects in effector CD8+ T cell expansion observed in Panx1 deficiency (**Figure S5d**). In contrast, P2RX7 deletion did not affect the *in vitro* activation of CD8^+^ T cells (**Figure S5e**).

**Figure 3.**
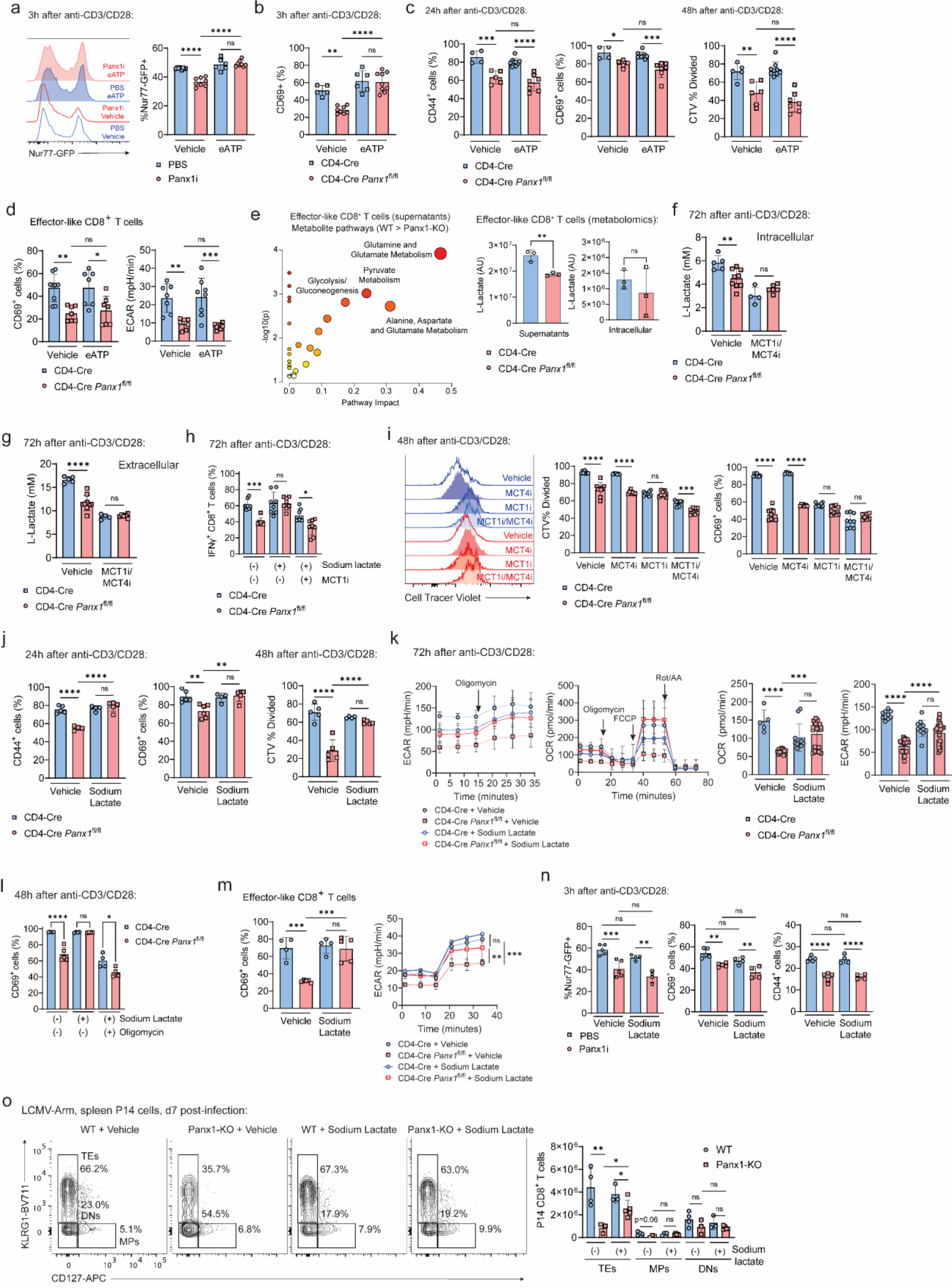
Panx1 controls the activation of effector CD8^+^ T cells through export of eATP and late extracellular lactate accumulation. (**a**) WT Nur77-GFP CD8^+^ T cells were activated in the presence of PBS or Panx1i, with the addition of vehicle or eATP. Nur77-GFP representative histograms (left) and average Nur77-GFP^+^ percentages (right) are shown. (**b-c**) CD4-Cre or CD4-Cre *Panx1*^fl/fl^ CD8^+^ T cells were activated *in vitro* (anti-CD3/CD28 + IL-2) for up to 48h, with the addition of vehicle or eATP at either the beginning of cultures (**b**), or at 20h after activation (**c**). (**b**) Average percentages of CD69^+^ cells at 3h after activation. (**c**) Average percentages of CD44^+^ and CD69^+^ cells at 24h after activation (left); average percentages of Cell Tracer Violet^-^ cells (CTV % divided) at 48h after activation (right). (**d**) CD4-Cre or CD4-Cre *Panx1*^fl/fl^ effector-like CD8^+^ T cells (with prolonged exposure to IL-2) were incubated in the presence or absence of eATP; average percentages of CD69^+^ cells and of ECAR values are shown. (**e**) WT (CD4-Cre) or Panx1-KO (CD4-Cre *Panx1*^fl/fl^) effector-like and memory-like CD8^+^ T cell cultures were harvested, and intracellular lysates and supernatants were submitted for untargeted metabolomics (GC-MS) analysis. Enrichment analysis showing pathways preferentially represented in the metabolites from the supernatants of WT effector-like CD8^+^ T cells (WT>Panx1-KO). Levels (arbitrary units – AU) of l-lactate in the supernatants (left) and intracellular lysates (right) of effector-like WT and Panx1-KO CD8^+^ T cells are shown below. (**f-g**) CD4-Cre or CD4-Cre *Panx1*^fl/fl^ CD8^+^ T cells were activated *in vitro* (anti-CD3/CD28 + IL-2) for 72h, and l-lactate measurements (mM) were done. (**f**) Intracellular lactate levels. (**g**) Extracellular lactate levels, in the presence or absence of inhibitors for MCT1 (SR13800) and MCT4 (VB124). (**h**) Average percentages of IFNγ^+^ CD8^+^ T cells at 72h after activation, with addition of sodium lactate +/- MCT1i. (**i**) Representative histograms (left), and average percentages of CTV % divided and CD69^+^ cells at 48h after activation, with addition of vehicle, MCT1i, MCT4i or MCT1/MCT4i. (**g-k**) CD4-Cre or CD4-Cre *Panx1*^fl/fl^ CD8^+^ T cells were activated *in vitro* (anti-CD3/CD28 + IL-2) for up to 72h, in the presence of the indicated metabolites or inhibitors. (**j**) Average percentages of CD44^+^ and CD69^+^ cells after 24h of activation (left), and of CTV % divided cells at 48h after activation (right), with the addition of vehicle or sodium lactate. (**k**) ECAR and OCR kinetics (left) and average baseline values (right) at 72h after activation, with the addition of vehicle or sodium lactate. (**l**) Average percentages of CD69^+^ cells at 48h after activation, with addition or sodium lactate +/-oligomycin. (**m**) CD4-Cre or CD4-Cre *Panx1*^fl/fl^ effector-like CD8^+^ T cells were incubated in the presence or absence of sodium lactate (right); average percentages of CD69^+^ cells and of ECAR values are shown. (**n**) WT Nur77-GFP CD8^+^ T cells were activated in the presence of PBS or Panx1i, with the addition of vehicle or sodium lactate; average percentages of Nur77-GFP^+^ cells (left), CD69^+^ cells (center), or CD44^+^ cells (right) are shown. (**o**) WT (CD4-Cre) or Panx1-KO (CD4-Cre *Panx1*^fl/fl^) P14 cells (CD45.2^+^) were transferred into LCMV-infected WT CD45.1^+^ mice. Some mice were treated with sodium lactate between days 1-3 post-infection, and spleen P14 cells were analyzed at day 7 post-infection. Flow cytometry plots showing expression of CD127 and KLRG1 (left) and the average numbers of TE, MP and DN P14 cells per spleen (right) are shown. (**a-d, f-o**) Data from 2-3 independent experiments, n=3-14 per experimental group. (**e**) Data from 3 biological replicates per experimental group (from n=3 mice per group). ns – not significant (p>0.05), *p<0.05, **p<0.01, ***p<0.001, ****p<0.0001, One-way ANOVA with Tukey’s post-test (**a-d, f-o**) Unpaired t-test (**e**), Two-way ANOVA with Bonferroni’s post-test (**m**).

In contrast, activation defects observed in Panx1-KO at 24h or later post-stimulation could not be restored by exogenous eATP (**Figure 3c**). Exogenous eATP also failed to rescue Panx1-KO defects in IL-2-induced effector-like CD8+ T cells (**Figure 3d**). These results suggest that other metabolites may be needed to restore the activation of Panx1-deficient CD8^+^ T cells. To investigate further, we used untargeted metabolomics to identify the intracellular and extracellular metabolites affected by Panx1-KO in effector-like and memory-like (IL-15-stimulated; further details below) CD8^+^ T cells (**Figure S5f-g**). We found >250 extracellular metabolites with decreased levels in effector-like Panx1-KO supernatants, and >300 in memory-like Panx1-KO supernatants (top 100 of each depicted in **Figure S5h**). Among the metabolites decreased in Panx1-KO effector-like CD8^+^ T cell supernatants, we found amino acids, fatty acids, and metabolites from the glycolysis pathway (**Figure 3e**). One of these metabolites, lactate, has recently been suggested to promote effector CD8^+^ T cell activation ^43,44, 45^, despite past contrary evidence ^46^. The accumulation of extracellular lactate is likely to be a consequence of decreased early activation in Panx1-KO CD8^+^ T cells, as intracellular lactate levels are also diminished (**Figure 3f**), and both intracellular and extracellular lactate levels in WT versus Panx1-KO CD8^+^ T cells are highly dependent on MCT1 and MCT4, the canonical lactate transporters (**Figures 3f-g**); mRNA expression of MCT4 (*Slc16a3*), but not MCT1 (*Slc16a1*), was affected in Panx1-KO activated CD8^+^ T cells (**Figure S5i**). Blockade of MCT1 and MCT4 significantly affected WT, but not Panx1-KO CD8^+^ T cell proliferation and expression of CD69 (**Figure 3h-i**), further suggesting that Panx1 promotes CD8^+^ T cell activation through indirect induction of extracellular lactate accumulation.

We then tested whether exogenous lactate would rescue the activation and proliferation of Panx1-KO CD8^+^ T cells. Sodium lactate (a pH-neutral lactate salt) significantly rescued Panx1-KO CD8^+^ T cell proliferation and activation (**Figure 3j**). Sodium lactate also significantly restored the glycolysis and mitochondrial respiration of activated Panx1-KO CD8^+^ T cells (**Figure 3k**).

The restoring effects of sodium lactate are abolished when the OXPHOS inhibitor Oligomycin is concomitantly present (**Figure 3l**). Effector-like defects in CD69 expression and glycolysis were restored by exogenous sodium lactate (**Figure 3m**). In contrast, early (3h post-stimulation) activation defects of Panx1-KO could not be restored by sodium lactate (**Figure 3n**). Finally, sodium lactate treatment significantly rescued the effector expansion of Panx1-KO P14 cells in LCMV-infected mice (**Figure 3o, Fig. S6**). These findings indicate that Panx1 promotes the early activation and full effector differentiation of CD8^+^ T cells through export of eATP and indirect control of lactate accumulation.

### Panx1 amplifies the consumption of carbon sources for full effector CD8^+^ T cell differentiation

We found that Panx1-KO CD8^+^ T cells have an overall defect in bioenergetic metabolism after activation. Indeed, the accumulation of several TCA cycle metabolites is diminished in effector-like CD8^+^ T cells (**Figure 4a**). Because consumption of carbon sources is necessary for the engagement of such pathways, we used carbon (^13^C) tracing metabolomics to measure the incorporation of glucose (**Figure 4b**) or lactate (**Figure 4c**). Compared to WT CD8^+^ T cells, Panx1-KO cells had decreased glucose incorporation of α-ketoglutarate, a rate-determining intermediate of the TCA cycle fundamental for energetic metabolism^47^. In contrast, lactate incorporation into citrate was decreased in Panx1-KO. Citrate, although a TCA cycle intermediate, can be shuttled out of the mitochondria and serve as a substrate for fatty acid synthesis^48^, which is crucial for the full activation of effector CD8^+^ T cells^49^.

**Figure 4.**
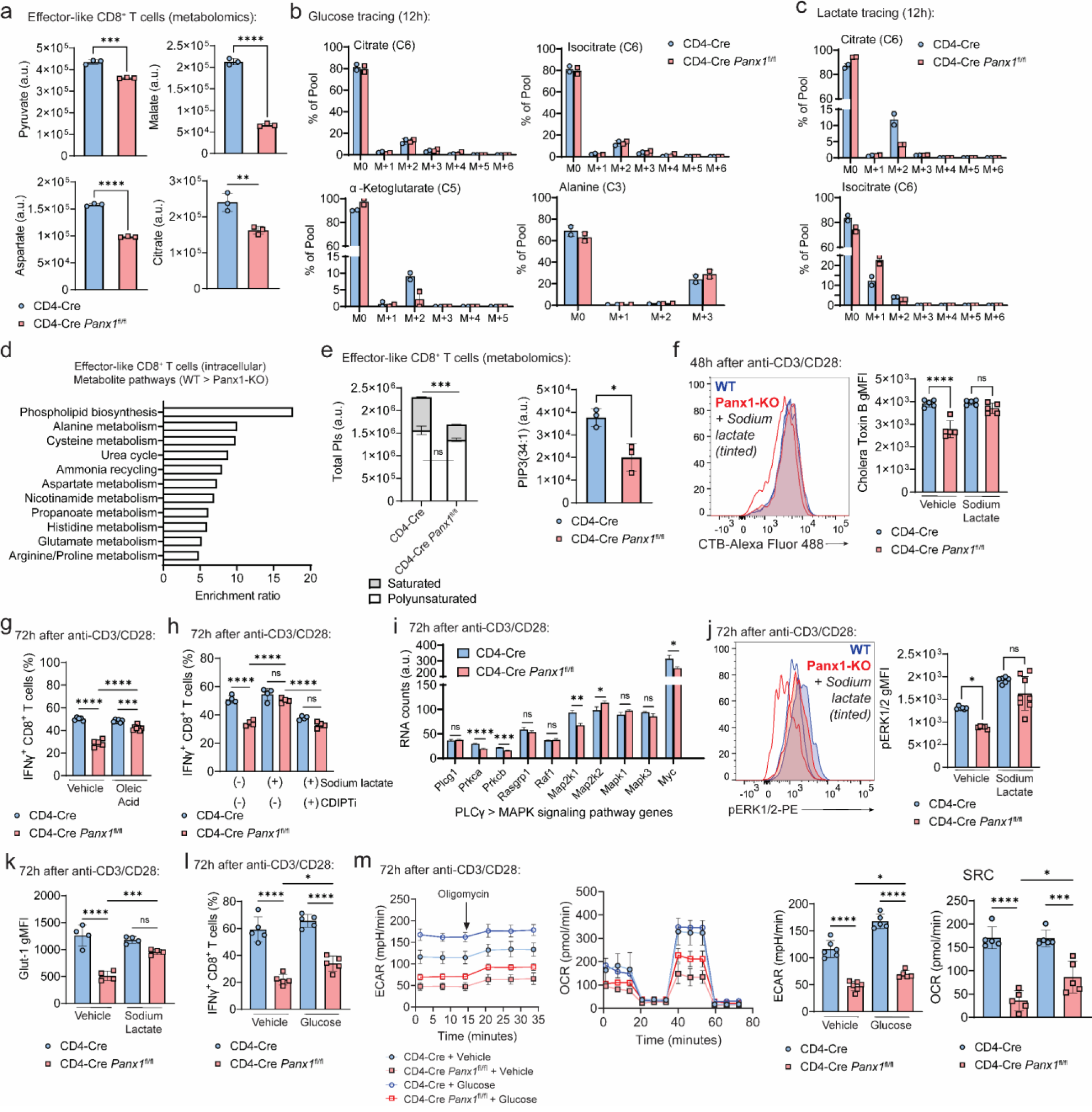
Panx1 promotes carbon source uptake and leads to maintenance of lipid rafts in activated CD8^+^ T cells. (**a**) Levels (arbitrary units – AU) of pyruvate, malate, aspartate, and citrate in the intracellular lysates of effector-like WT (CD4-Cre) or Panx1-KO (CD4-Cre *Panx1*^fl/fl^) CD8^+^ T cells from untargeted metabolomics are shown. (**b-c**) CD4-Cre and CD4-Cre *Panx1*^fl/fl^ CD8^+^ T cells were activated *in vitro* (anti-CD3/CD28 + IL-2) for 72h; in the last 12h, either ^13^C-Glucose (**b**) or ^13^C-Lactate (**c**) were substituted into the cultures for metabolite carbon tracing. (**b**) Percentages of ^13^C-Glucose incorporation into citrate, isocitrate, a-ketoglutarate, and alanine. (**c**) Percentages of ^13^C-Lactate incorporation into citrate and isocitrate. (**d**) Top enriched intracellular pathways in WT effector-like CD8^+^ T cells (compared to Panx1-KO), depicted from the untargeted metabolomics from (**a**). (**e**) Levels (AU) of total PIs in CD4-Cre versus CD4-Cre *Panx1*^fl/fl^ effector-like CD8^+^ T cells; polyunsaturated (2 or more double bonds) or saturated (1 or less double bonds) PIs are depicted in white and grey, respectively. In the right, levels (AU) of PIP(34:1) in CD4-Cre versus CD4-Cre *Panx1*^fl/fl^ effector-like CD8^+^ T cells. (**f**) CD4-Cre and CD4-Cre *Panx1*^fl/fl^ CD8^+^ T cells were activated *in vitro* (anti-CD3/CD28 + IL-2) in the presence or not of Sodium Lactate. Representative histograms (left) and average gMFI values (right) for Cholera Toxin B are shown. (**g**) CD4-Cre and CD4-Cre *Panx1*^fl/fl^ CD8^+^ T cells were activated *in vitro* (anti-CD3/CD28 + IL-2) in the presence or not of BSA-conjugated Oleic Acid. Average percentages of IFNγ^+^ CD8^+^ T cells are shown. (**h**) CD4-Cre and CD4-Cre *Panx1*^fl/fl^ CD8^+^ T cells were activated *in vitro* (anti-CD3/CD28 + IL-2) in the presence or not of Sodium Lactate and/or the CDIPT inhibitor Inostamycin (CDIPTi). Average percentages of IFNγ^+^ CD8^+^ T cells are shown. (**i**) RNA expression levels (AU) of the indicated PLCγ-MAPK signaling pathway genes in CD4-Cre and CD4-Cre *Panx1*^fl/fl^ CD8^+^ T cells after 72h of activation. (**j**) CD4-Cre and CD4-Cre *Panx1*^fl/fl^ CD8^+^ T cells were activated *in vitro* (anti-CD3/CD28 + IL-2) in the presence or not of Sodium Lactate. Representative histograms (left) and average gMFI values (right) for phosphorylated ERK1/2 (pERK1/2) are shown. (**k**) CD4-Cre and CD4-Cre *Panx1*^fl/fl^ CD8^+^ T cells were activated *in vitro* (anti-CD3/CD28 + IL-2) in the presence or not of Sodium Lactate. Average gMFI values for Glut1 are shown. (**l-m**) CD4-Cre and CD4-Cre *Panx1*^fl/fl^ CD8^+^ T cells were activated *in vitro* (anti-CD3/CD28 + IL-2) in the presence or not of additional Glucose. (**l**) Average percentages of IFNγ^+^ CD8^+^ T cells are shown. (**m**) ECAR kinetics, OCR kinetics, baseline ECAR and SRC OCR values are respectively shown. (**a-e, i**) Data from 2-3 biological replicates per experimental group (from n=3 mice per group). (**f-h, j-m**) Data from 2-3 independent experiments, n=4-7 per experimental group. ns – not significant (p>0.05), *p<0.05, **p<0.01, ***p<0.001, ****p<0.0001, Unpaired t-test (**a, e, i**), One-way ANOVA with Tukey’s post-test (**f-h, j-m**).

In screenings to identify additional metabolites (among the ones depicted in **Figure S5h**) that rescue Panx1-KO CD8^+^ T cell activation (**Figure S5j**), among the few other candidates found was Hydroxylauric Acid, a medium-chain fatty acid ^50^. Medium-chain fatty acids are used by CD8^+^ T cells for fatty acid biosynthesis. Intracellular phospholipid biosynthesis metabolites were sharply decreased in Panx1-KO effector-like CD8^+^ T cells (**Figure 4d**), including phosphatidylinositol (PIs) (**Figures 4e, S5k**). PIs compose the immunological synapse ^51^, and saturated (i.e., with less than 2 double bonds) phosphatidylinositol phosphates (PIPs) compose lipid rafts found during late activation, to sustain effector function ^49^. Panx1-KO effector-like CD8^+^ T cells had specific defects in saturated PIs (**Figures 4e, S5k**) and decreased lipid raft formation as depicted by Cholera Toxin B (CTB) staining (**Figure 4f**) – which can be restored by sodium lactate, in an MCT-1-dependent way (**Figure S5l**). Additionally, oleic acid, a PI biosynthesis-forming medium-chain fatty acid ^49^, had decreased intracellular levels in Panx1-KO CD8^+^ T cells (**Figure S5m**) – possibly as a consequence from impaired initial activation and engagement of bioenergetic metabolism. Exogenous oleic acid partially (but not completely) restored Panx1-KO CD8^+^ T cell activation (**Figure 4g**), suggesting that Panx1-mediated accumulation of this fatty acid plays a role in effector CD8^+^ T cell activation. The restoring effects of sodium lactate for Panx1-KO CD8^+^ T cell activation were abolished when an inhibitor of CDIPT, the enzyme that catalyzes the biosynthesis of saturated fatty acids, is present (**Figure 4h, S5n**). Saturated PI biosynthesis leads to activation of the MAP-kinase (MAPK) signaling pathway to sustain CD8^+^ T cell activation and effector function ^49^. RNA expression of several MAPK pathway genes is decreased in Panx1-KO effector-like CD8^+^ T cells (**Figure 4i**), and phosphorylated ERK1/2 levels are diminished in Panx1-KO CD8^+^ T cells – which is restored in the presence of sodium lactate (**Figure 4j**). Therefore, the Panx1-induced effects of sodium lactate are associated with saturated PI generation and formation of lipid rafts.

Because of the Panx1 effects on glycolysis and the Panx1-dependent incorporation of glucose into the TCA cycle, we investigated how Panx1-mediated activation could be dependent of exogenous glucose. Glut1 expression is diminished in Panx1-KO *in vitro*-activated CD8^+^ T cells, which was restored by exogenous sodium lactate (**Figure 4k**). Exogenous glucose addition to *in vitro* cultures led to minor, but statistically significant increases in IFN-γ production, glycolysis, and mitochondrial respiration (**Figures 4l-m**). These results suggest that Panx1 promotes the full effector differentiation of CD8^+^ T cells through an increase in the consumption of carbon sources.

### Panx1 is crucial for the survival of memory CD8^+^ T cells

Panx1 mRNA is also found in memory CD8^+^ T cell subsets (**Figure 1a, S2a**). We also investigated how Panx1 affects memory CD8^+^ T cell establishment (gating strategies in **Figure S1**). T cell-specific Panx1-KO resulted in significantly decreased numbers of circulating memory (**Figure 5a**) and small intestine intraepithelial (SI-IEL) tissue-resident memory (T_RM_; **Figure 5b**) gp33-specific CD8^+^ T cells. Similar defects were observed in 1:1 WT:Panx1-KO mixed bone marrow chimeras, (**Figures 5c-d**). To test whether memory CD8^+^ T cell survival is specifically regulated by Panx1, we used a tamoxifen-inducible KO system (**Figure 5e**). Tamoxifen-induced Panx1 ablation during the effector stage affected both the accumulation of effector CD8^+^ T cells and the establishment of memory CD8^+^ T cells (**Figure S7**). Panx1-KO after memory establishment resulted in a steady decline of CD8^+^ T cell numbers (**Figure 5f**). The decrease in memory CD8^+^ T cell numbers with late Panx1 ablation was true for all subsets of circulating memory, although the most pronounced decline was in central memory (T_CM_) cells (**Figures 5g-h**). Late Panx1-KO also led to a decrease in T_RM_ cells, except for the SI-IEL compartment (**Figure 5i**). These results highlight a crucial role for Panx1 specifically for the long-term survival of memory CD8^+^ T cells.

**Figure 5.**
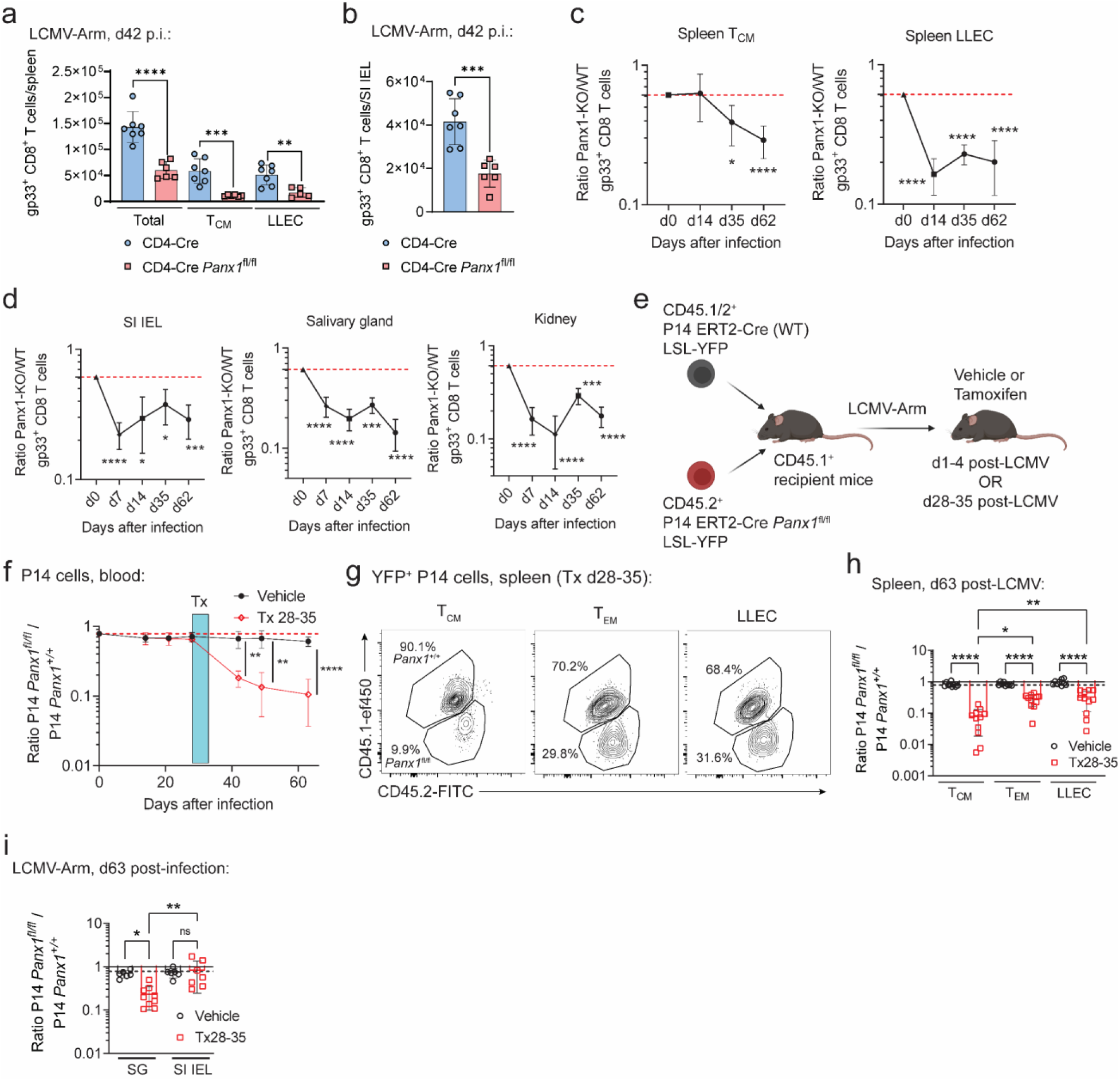
Panx1 promotes the long-term establishment of memory CD8^+^ T cells. (**a-b**) CD4-Cre or CD4-Cre *Panx1*^fl/fl^ mice were infected with LCMV-Arm and gp33^+^ CD8^+^ T cells were tracked over time. Numbers of total, T_CM_ and LLEC spleen gp33^+^ CD8^+^ T cells at day 42 post-infection are shown in (**a**), while the numbers of SI IEL gp33^+^ CD8^+^ T cells at day 42 post-infection are shown in (**b**). (**c-d**) A 1:1 mix of bone marrow cells from CD4-Cre (WT; CD45.1^+^) and CD4-Cre *Panx1*^fl/fl^ (Panx1-KO; CD45.2^+^) mice was transferred into lethally irradiated C57BL/6 mice (CD45.1/2^+^). After two months, BM chimeric mice were infected with LCMV-Arm. The red dotted lines show the Panx1-KO:WT ratios after two months of reconstitution. Panx1-KO/WT ratios of T_CM_ and LLEC gp33^+^ spleen CD8^+^ T cells over time post-infection are shown in (**c**). In (**d**), the Panx1-KO/WT ratios of SI IEL, Salivary gland, and kidney T_RM_ gp33^+^ spleen CD8^+^ T cells over time post-infection are shown. (**e-i**) Recipient C57BL/6 mice (CD45.1^+^) were adoptively transferred with a 1:1 mix of P14 ERT2-Cre LSL-YFP *Panx1*^+/+^ (WT; CD45.1/2^+^) and P14 ERT2-Cre LSL-YFP *Panx1*^fl/fl^ (CD45.2^+^) cells, then infected with LCMV-Arm. Infected mice were treated with vehicle or tamoxifen (Tx) at the indicated time intervals. (**e**) Experimental design. (**f**) *Panx1*^fl/fl^/*Panx1*^+/+^ P14 cell ratios in the blood over time post-infection, in mice treated with vehicle or Tx between days 28-35 (d28-35) post-infection. Treatment periods is indicated by the blue shaded box. (**g**) Representative flow cytometry plots showing the distribution of spleen T_CM_, T_EM_ and LLEC *Panx1*^+/+^ and *Panx1*^fl/fl^ P14 cells from mice treated with Tx d28-35 at 63d post-infection. (**h**) *Panx1*^fl/fl^/*Panx1*^+/+^ ratios of spleen T_CM_, T_EM_ and LLEC P14 cells from mice treated with vehicle or Tx d28-35, at 63d post-infection. (**i**) *Panx1*^fl/fl^/*Panx1*^+/+^ P14 cell ratios in SG and SI IEL at day 63 post-infection, in mice treated with Tx between days 28-35. (**a-d, f-i**) Data from 3 independent experiments, n=6-12 per time point per experimental group. ns – not significant (p>0.05), *p<0.05, **p<0.01, ***p<0.001, ****p<0.0001, One-way ANOVA with Tukey’s post-test (**a, h-i**), Unpaired t-test (**b**), Two-way ANOVA with Bonferroni’s post-test (**c-d, f**).

### Panx1 promotes memory CD8^+^ T cell survival through the eATP receptor P2RX7 and the AMPK metabolic pathway

Panx1 is canonically described as an ATP exporter and has been linked with activation of the eATP receptor P2RX7 ^20,23–27,32,52^. We had previously found a crucial role for P2RX7 in the induction of memory CD8^+^ T cells^32,34,37^. To test if Panx1 regulates memory CD8^+^ T cells through P2RX7, we used CRISPR-Cas9 ^53^ to induce Panx1-KO in WT and P2RX7-KO P14 cells, thus generating double-knockout (DKO) CD8^+^ T cells (**Figure 6a**). Upon adoptive transfer and LCMV infection, we observed similar P14 T_CM_ cell numbers between Panx1-KO, P2RX7-KO and DKO groups – all lower than WT P14 T_CM_ cells (**Figure 6b**). These data suggest that Panx1 may regulate T_CM_ cell establishment through P2RX7. LLEC P14 cell numbers were further diminished in DKO cells if compared to either single-KO, suggesting a non-redundant role of Panx1 and P2RX7 for this subset (**Figure 6b**, right). P2RX7 promotes memory CD8^+^ T cell differentiation through promotion of mitochondrial respiration^32^. Panx1-KO mirrored this phenotype, with sorted MP cells displaying reduced mitochondrial respiration compared to WT TE cells (**Figure 6c**). To better understand how Panx1 regulates the mitochondrial metabolism of memory-phenotype CD8^+^ T cells, we used prolonged IL-15 stimulation of *in vitro*-activated cells to generate memory-like conditions^41^ (**Figure 6d**). Panx1-KO memory-like CD8^+^ T cells displayed reduced mitochondrial respiration and no defects on glycolysis – leading to increased ECAR/OCR ratios (**Figure 6e**). The Panx1-KO memory-like defects in mitochondrial respiration and homeostasis were significantly restored by exogenous eATP, and this rescue effect was blocked by the P2RX7 inhibitor A-438079 (**Figure 6f-g**). Panx1 also induced the extracellular accumulation of lactate in memory-like CD8^+^ T cells (**Figure S5g**), but sodium lactate could not restore the mitochondrial defects of Panx1-KO memory-like CD8^+^ T cells (**Figure 6h**). These data suggest Panx1 promotes the mitochondrial fitness of memory-phenotype CD8^+^ T cells through induction of eATP accumulation and activation of P2RX7.

**Figure 6.**
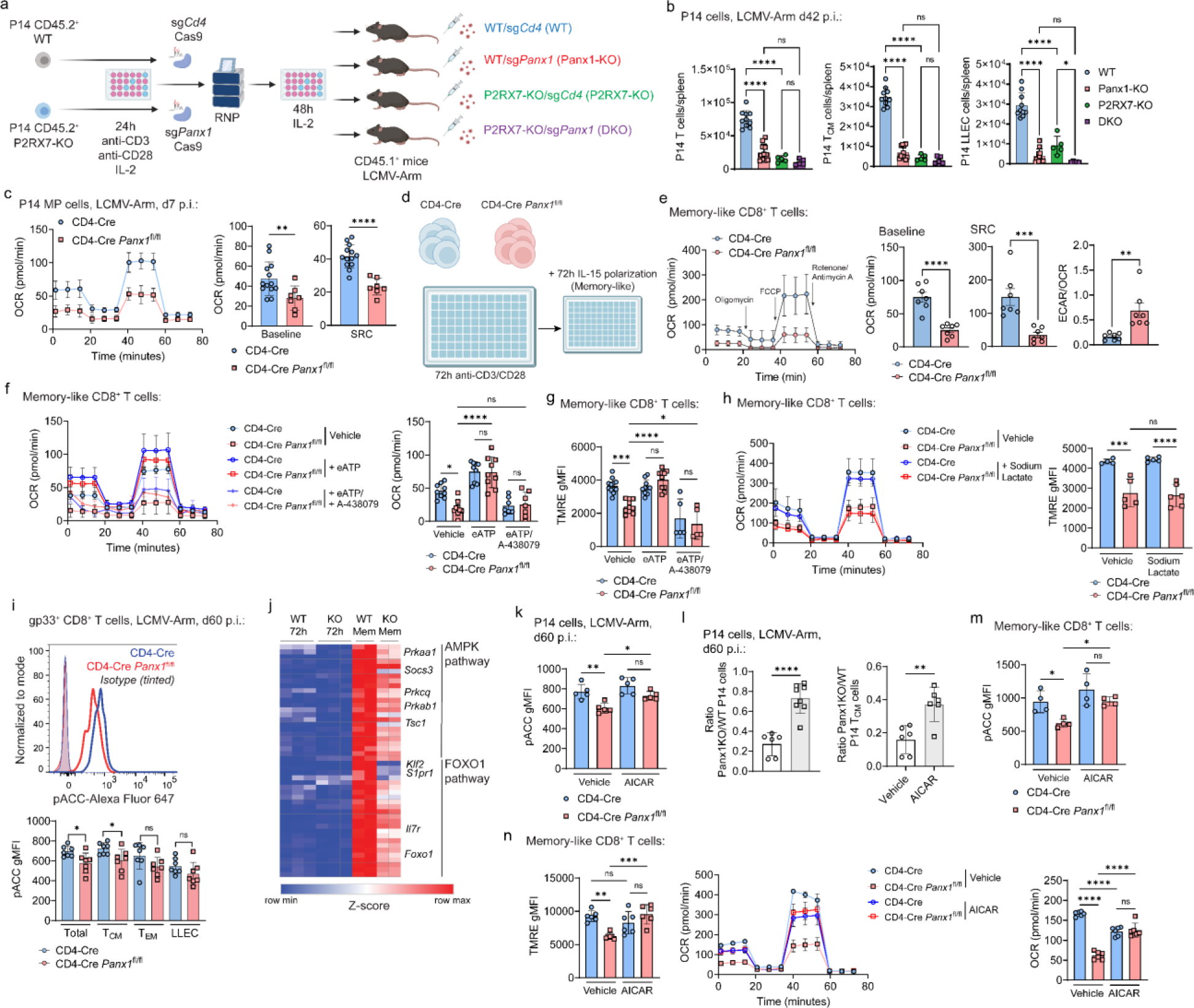
Panx1 promotes memory CD8^+^ T cell maintenance through eATP export and activation of the AMPK pathway. (**a-b**) WT (CD4-Cre) or P2RX7-KO (CD4-Cre *P2rx7*^fl/fl^) P14 cells (CD45.2^+^) were *in vitro* activated for 24h, then were electroporated with the Cas9 protein combined with sgRNAs for *Cd4* (sg*Cd4*; control) or *Panx1* (sg*Panx1*; Panx1-KO). After 48h of resting with IL-2, P14 cells were transferred into recipient mice (CD45.1^+^) infected with LCMV-Arm. (**a**) Experimental design. (**b**) Numbers of total spleen P14 cells (left), spleen T_CM_ P14 cells (middle) and spleen LLEC P14 cells (right) at day 42 post-infection. (**c**) CD4-Cre and CD4-Cre *Panx1*^fl/fl^ P14 cells were transferred into C57BL/6 mice infected with LCMV-Arm. At day 7 post-infection, MP P14 cells were sorted. OCR kinetics (left), average baseline OCR values (middle) and average SRC OCR values (right) are shown. (**d-h**) CD4-Cre or CD4-Cre *Panx1*^fl/fl^ CD8^+^ T cells were activated *in vitro* (anti-CD3/CD28 + IL-2) for 72h, then further polarized into memory-like cells with IL-15 for additional 72h. (**d**) Experimental design. (**e**) In the left, OCR kinetics for both groups are shown. In the right, average baseline OCR values, average SRC OCR values, and average ECAR/OCR ratios are shown. (**f**) OCR kinetics (left) and average baseline OCR values (right) are shown for memory-like cells with additional treatment with eATP +/- A-438079. (**g**) Average TMRE gMFI values for memory-like cells with additional treatment with eATP +/- A-438079. (**h**) OCR kinetics (left) and average TMRE gMFI values for memory-like cells with additional treatment with sodium lactate. (**i**) CD4-Cre or CD4-Cre *Panx1*^fl/fl^ mice were infected with LCMV-Arm and analyzed at 60 days post-infection. Histograms showing pACC staining (top) and average pACC gMFI values (bottom) are shown for memory gp33^+^ CD8^+^ T cells. (**j**) Heatmap showing expression of genes from the AMPK and FOXO1 pathways in memory-like CD8^+^ T cells; expression in CD8^+^ T cells (with 72h post-activation) are shown in comparison. (**k-l**) CD4-Cre (WT; CD45.1/2^+^) or CD4-Cre *Panx1*^fl/fl^ (Panx1-KO; CD45.2^+^) P14 cells were transferred into WT (CD45.1^+^) mice infected with LCMV-Arm; between days 1-7 post-infection, mice were treated with AICAR. (**k**) pACC expression in P14 cells. (**l**) Panx1-KO/WT P14 cell ratios for spleen total (top) and T_CM_ (bottom) cells are shown. (**m-n**) CD4-Cre or CD4-Cre *Panx1*^fl/fl^ CD8^+^ T cells were activated *in vitro* (anti-CD3/CD28 + IL-2) then polarized into memory-like cells, with additional treatment with vehicle or AICAR. (**m**) pACC expression in CD8^+^ T cells. (**n**) Average TMRE gMFI values (left), OCR kinetics (middle), and average baseline OCR values (right) are shown. (**b-c, e-i, k-n**) Data from 2-3 independent experiments, n=4-13. (**j**) Data from 2-3 biological replicates per experimental group (from n=5-8 mice per group). ns – not significant (p>0.05), *p<0.05, **p<0.01, ***p<0.001, ****p<0.0001, One-way ANOVA with Tukey’s post-test (**b, f-i, m-n**), Unpaired t-test (**c, e, k-l**).

We have previously established that P2RX7 promotes memory CD8^+^ T cells partly through activation of the AMPK pathway ^32^. AMPK can be triggered, among other signals, by increased AMP-to-ATP ratios ^54,55^. Since Panx1 blockade promoted increased intracellular ATP (**Figure S2b**), we monitored activation of the AMPK pathway in Panx1-KO memory CD8^+^ T cells. Panx1-KO led to decreased phosphorylation of the downstream AMPK target Acetyl-CoA-Carboxylase (pACC) in memory CD8^+^ T cells, especially T_CM_ cells (**Figure 6i**). In addition, Panx1-KO memory-like CD8^+^ T cells had reduced mRNA expression of several genes associated with the AMPK pathway (**Figure 6j**). This suggests a connection between Panx1 promotion of memory CD8^+^ T cells and activation of the AMPK pathway. To test whether Panx1 promotes memory CD8^+^ T cells through AMPK, we co-transferred WT and Panx1-KO P14 cells into LCMV-infected mice that were treated with AICAR, an AMPK pathway agonist. The Panx1-KO/WT P14 cell ratios at the memory phase were significantly increased in AICAR-treated mice (**Figure 6k-l**), suggesting a preferential rescue of Panx1-KO memory CD8^+^ T cell generation by forced AMPK activation. Moreover, Panx1-KO memory-like CD8^+^ T cells cultured in the presence of AICAR displayed increased mitochondrial function (**Figure 6m-n**). Our results indicate that Panx1 promotes the mitochondrial fitness and long-term establishment of memory CD8^+^ T cells through accumulation of eATP, induction of P2RX7 activation, and promotion of the AMPK signaling pathway.

### Panx1 deficiency hinders tumor control or graft-versus-host disease induced by CD8^+^ T cells

Given the importance of CD8^+^ T cell responses in controlling solid tumors ^56–58^, we assessed the role of Panx1 in CD8^+^ T cell-mediated melanoma control. To circumvent any non-CD8+ T cell-specific effects of Panx1, we used an adoptive cell therapy (ACT) model previously used by our group^59,60^. *In vitro*-activated Panx1-KO P14 cells showed defects in limiting B16-gp33 tumor growth and fail to promote survival of B16-gp33 tumor-bearing mice compared to activated WT P14 cells (**Figures 7a-c**). We next evaluated the numbers and phenotype of adoptively transferred WT and Panx1-KO P14 cells in tumor-bearing mice through co-transfer experiments. We observed a significant underrepresentation of Panx1-KO P14 cells infiltrating B16 tumors, with no differences detected in draining lymph nodes or spleen (**Figure 7d**). Among tumor-infiltrating P14 cells, Panx1-KO cells exhibited increased signs of mitochondrial dysfunction such as depolarized mitochondria, as well as decreased levels of mitochondrial mass or membrane potential (**Figure 7e**). These data suggest that expression of Panx1 is important for CD8^+^ T cells to infiltrate tumors, maintain mitochondrial homeostasis, and perform optimal antitumor function.

**Figure 7.**
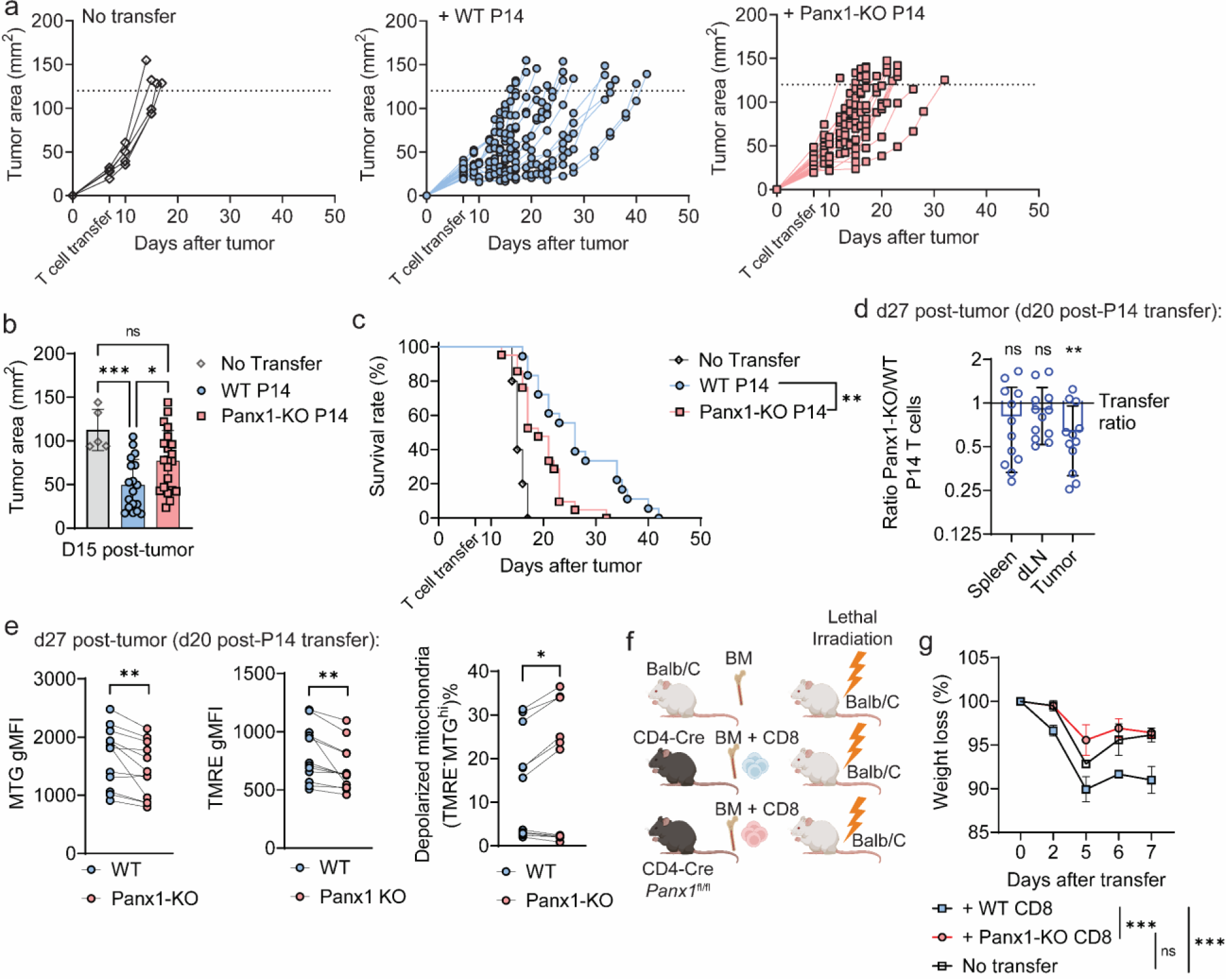
Cell-specific Panx1 promotes antitumor CD8^+^ T cell responses and graft-versus-host disease *in vivo*. (**a-e**) WT (CD4-Cre) and Panx1-KO (CD4-Cre *Panx1*^fl/fl^) P14 cells were activated *in vitro* (α-CD3, α-CD28, IL-2 and IL-12) for 72h. After this period, P14 cells were transferred individually (**a-c**) or co-transferred (**d-e**) into B16.gp33 tumor-bearing mice. The tumor sizes and survival of individually transferred mice were tracked over time post-tumor inoculation; the numbers and phenotype of transferred P14 cells were assessed at 20 days post-tumor inoculation in co-transferred mice. (**a**) Individual tumor growth values over time. (**b**) Average tumor areas at day 14 post-tumor inoculation are shown. (**c**) Survival curves of individually transferred mice over time. (**d**) Average Panx1-KO/WT P14 cell number ratios in the indicated organs. (**e**) Average values for Mito Green (MTG) gMFI (left), TMRE gMFI (middle) and percent of P14 cells with depolarized mitochondria (MTG^hi^TMRE^-^; right). (**f-g**) In other experiments, BALB/c mice were lethally irradiated and reconstituted with either BALB/c BM cells (No transfer), BM + spleen CD8^+^ T cells from CD4-Cre mice (+ WT CD8) and BM + spleen CD8^+^ T cells from CD4-Cre *Panx1*^fl/fl^ mice (+ Panx1-KO CD8). (**f**) Experimental design. (**g**) Average survival values over time after cell transfer. (**a-g**) Data from 2-3 independent experiments, n=5-20 per experimental group. ns – not significant (p>0.05), *p<0.05, **p<0.01, ***p<0.001, Kaplan-Meier Survival curve analysis (**e**), Unpaired t-test (**g-f**), One-way ANOVA with Tukey’s post-test (**b**), Two-way ANOVA with Bonferroni’s post-test (**g**).

Finally, we tested the biological relevance of Panx1 for CD8^+^ T cell disease-inducing responses using an experimental model of graft-versus-host disease (GVHD). This model relies on the adoptive transfer of naïve CD8^+^ T cells from C57BL/6 mice into MHC-mismatched BALB/c mice^61^ (**Figure 7f**). Transfer of WT, but not CD4-Cre *Panx1*^fl/fl^ CD8^+^ T cells led to a precipitous weight loss in recipient BALB/c mice (**Figure 7g**). These data indicate that Panx1 promotes CD8^+^ T cell responses that can cause deleterious effects, suggesting the potential utility of Panx1 inhibition as a therapeutic target in these conditions.

## Discussion

Sensing of extracellular metabolites promotes CD8^+^ T cell metabolism, homeostasis, and function ^62–64^. The sources of these metabolites and how they affect CD8^+^ T cell intracellular pathways are still not fully understood. Our work suggests that, through the action of Panx1, CD8^+^ T cells utilize eATP for their own sensing, promoting CD8^+^ T cell responses at the effector and at the memory stage. Additionally, we found that Panx1, besides its well-known role as an eATP exporter^20^, can regulate CD8^+^ T cell responses through indirect accumulation of extracellular lactate. Both effector and memory-phenotype CD8^+^ T cells induce eATP and lactate extracellular accumulation via Panx1, ruling out differential metabolite export as a reason why Panx1 controls effector and memory CD8^+^ T cell establishment differently. Rather, the outcome of the Panx1 role is dictated by the signaling pathways and specific requirements for effector or memory CD8^+^ T cells. In future studies, we will investigate whether this phenomenon is generalizable for all metabolites exported via Panx1, or if certain molecules are exported differentially in effector versus memory CD8^+^ T cells.

We found a fundamental role of Panx1 in promoting effector CD8^+^ T cell responses. Consistent with previous reports, Panx1 controls the early activation of CD8^+^ T cells through eATP export, in a TCR priming and P2RX4-dependent way^42,65^. Unexpectedly, the role of Panx1 for the late stages of CD8^+^ T cell activation was not rescued by eATP. Instead, our data suggest that Panx1-mediated accumulation of extracellular lactate, likely as a byproduct of early glycolytic metabolism^66,67^, promotes the complete effector CD8^+^ T cell differentiation. While lactate has been considered immunosuppressive due to its coupling with acidosis^46,68^, lactate-induced acidosis is primarily observed in hypoxic conditions, such as in the tumor microenvironment ^69^. During T cell priming, lactate is present in a pH neutral condition, which is well-tolerated by cells^70^. Our results indicate that Panx1-induced extracellular lactate can be recycled by CD8^+^ T cells via MCT-1, leading to saturated PI biosynthesis, lipid raft formation, induction of MAPK pathway activation, and reinforcement of the effector CD8^+^ T cell program. Our work provides additional evidence that carbon sources induce the generation of optimal lipid raft structures for sustained CD8^+^ T cell activation ^49^. Despite a minor role for glucose in our model, glucose-independent sources such as lactate can be relevant carbon sources for CD8^+^ T cell activation and effector differentiation ^43,44,66^. Panx1, therefore, may serve as a “link” between these two concepts by indirectly inducing the accumulation of an extracellular lactate pool, which in turn is utilized by activated CD8^+^ T cells to sustain its effector differentiation.

Panx1 also promotes the survival of memory CD8^+^ T cells. Our data suggest that Panx1 favors the homeostasis of memory CD8^+^ T cells through the export of eATP, which is, in turn, sensed by P2RX7. These results are consistent with previous reports on *in vitro*-stimulated T cells ^31,52^ and support our speculation that, during the memory phase, eATP levels are tightly regulated around the pericellular space of memory CD8^+^ T cells ^32,71^. This process may seem, at first, energetically futile: the export of ATP would lead to a decrease in intracellular levels and subsequent lower availability for energy-demanding processes. However, there are potential advantages to Panx1-induced eATP export. For example, it would ensure constant engagement of P2RX7 to sustain Ca^2+^ influx and subsequent mitochondrial health ^72^. Indeed, Panx1-KO memory-phenotype CD8^+^ T cells display defective mitochondrial respiration. The exceptions were T_RM_ cells forming in tissues where potential alternative eATP sources are present: the SI-IEL, where eATP can be microbiota-derived ^37,73^, and the influenza-infected lung, where release of eATP by tissue damage may occur. Moreover, Panx1 export of ATP also impacts its intracellular levels^32^ and we have previously speculated the possible consequences of Panx1-smediated intracellular ATP decrease for CD8^+^ T cells ^71^. Promotion of the AMPK pathway due to decreased intracellular ATP in an LKB-dependent pathway favors memory CD8^+^ T cell homeostasis ^74^, which we have shown to rely on P2RX7 ^32^. Here, we found that Panx1 promotes memory CD8^+^ T cells through AMPK induction. Panx1-mediated intracellular metabolite alterations could also have effects on lactate. Excess intracellular lactate can hinder T cell activation ^75^, therefore Panx1-mediated lactate export can prevent this shutdown. We will investigate this possibility in the future, which would suggest Panx1 also acts as a “metabolite escape valve” for effector or memory CD8^+^ T cells.

CD8^+^ T cell-specific Panx1 also promotes melanoma control. At first, these results conflict with human cancer data, which shows high *PANX1* gene expression linked with poor prognosis in multiple tumors ^76^, including melanoma ^77^. Our data, however, focus on the CD8^+^ T cell-specific roles of Panx1. Indeed, a recent report has shown that *PANX1* expression and CD8^+^ T cell abundance is linked to better prognosis against stage III colorectal cancer^78^. Our ACT experiments suggest a model where Panx1 expression specifically in CD8^+^ T cells promote efficient tumor control. Because of the mitochondrial defects observed in intratumoral CD8^+^ T cells lacking Panx1, we speculate that, in the context of melanoma, the dominant role of Panx1 is to promote the longevity and homeostasis of tumor CD8^+^ T cells. Additional studies will be necessary to further understand this mechanism.

Here, we demonstrated that CD8^+^ T cells need Panx1 channels for their proper activation, expansion, and survival into memory phase. Beyond the classical nucleotide exporter role of Panx1, our results unveiled an unexpected link between expression of Panx1 and extracellular accumulation of lactate – which is necessary for the effector differentiation of CD8^+^ T cells.

Defining how Panx1 acts as an amplifier of CD8^+^ T cell responses will open new directions on our understanding of how adaptive immune cells interact with their surrounding milieu in response to disease. Our data suggests that, at least in part, CD8^+^ T cells can use Panx1 to generate a favorable extracellular microenvironment by recycling products of their own intracellular metabolic pathways.

## Acknowledgments

We thank the Borges da Silva and Lancaster labs (Mayo Clinic Arizona) for their intellectual input. We are thankful to the animal facilities at Mayo Clinic Arizona and at the University of Minnesota for technical assistance and mouse husbandry; T. Brehm-Gibson, C. Viso, I. Silva-Junior and E. Martinez for cell sorting and Flow Cytometry Core Facility maintenance at Mayo Clinic Arizona; and M. Petterson for technical assistance at the Mayo Clinic Rochester Metabolomics Core. We thank K. Ravichandran (U-Virginia, now Washington University) for providing the *Panx1*^fl/fl^ mice. We thank G. Hasko (Rutgers University) and Matyas Sandor (U-Wisconsin) for providing the *P2rx7*^fl/fl^ mice. BGI Americas provided bioinformatic support and resources that helped achieve the results reported within this paper. This work was supported by an individual predoctoral F30 fellowship from the National Institutes of Health (NIH) (F30CA250321) to K.M.W.; NIH grants R01AI038903 and R01AI145147 to S.C.J.; Mayo Clinic Center for Biomedical Discovery Pilot Grant-2021, and NIH grants K99/R00A139381 and R01A170649 to H.BdS.

## Author contributions

Conceptualization, T.V-K., H.BdS.; Methodology, T.V-K., H.BdS.; Formal Analysis, T.V-K., A.B., M.G-P., B.G.M., K.M.W., C.L.L., S.C.J. and H.BdS.; Investigation, T.V-K., A.B., M.G-P., B.G.M., C.L.S., K.M.W., M.H.Z., S.vD., I.S-C., A.S.B., C.L.L., C.P., and H.BdS.; Resources, N.L.T., S.C.J. and H.BdS.; Data Curation, T.V-K., A.B., M.G-P., B.G.M. and H.BdS.; Writing – Original Draft, T.V-K. and H.BdS.; Writing – Review & Editing, T.V-K., A.B., M.G-P., B.G.M., C.L.S., K.M.W., S.C.J, and H.BdS.; Supervision, H.BdS.; Project Administration, H.BdS.; Funding Acquisition, S.C.J. and H.BdS.

## Inclusion and Diversity

One or more of the authors of this paper self-identifies as an underrepresented ethnic minority in their field of research or within their geographical location. One or more of the authors of this paper self-identifies as a member of the LGBTQIA+ community.

## Materials and Methods

### Lead contact

Further information and requests for resources and reagents should be directed to and will be fulfilled by the lead contact, Henrique Borges da Silva.

### Materials availability

No new unique reagents were generated in this study.

### Data and code availability

- The data that support the findings of this study are available from the corresponding authors on request.
- All sequencing reads generated in this study and processed RNA expression matrices are deposited in NCBI’s GEO and are publicly available as of the date of publication. The accession numbers GSE217292 and GSE218643 are correspondent to the RNAseq analyses generated in this paper.
- Any additional information required to reanalyze the data reported in this paper is available from the lead contact upon request.

### Mice

Male and female 6-to 8-week-old adult BALB/c, C57BL/6 (B6) and B6.SJL (expressing the CD45.1 allele) mice were purchased from Jackson and Charles River and were allowed to acclimate to our housing facilities for at least one week. CD4-Cre, Nur77-GFP, ERT2-Cre and R26-EYFP (LSL-YFP) mice were obtained from Jackson Laboratories. *P2rx7*^fl/fl^ mice were obtained from Drs. Gyorgy Hasko (Rutgers University) and Matyas Sandor (U-Wisconsin). *Panx1*^fl/fl^ mice were obtained from Dr. Kodi Ravichandran (U-Virginia; now Washington University). LCMV-D^b^ GP33-specific TCR transgenic P14 mice were bred in-house at the University of Minnesota and then transferred to Mayo Clinic Arizona. P14 mice were fully backcrossed to B6, CD4-Cre *P2rx7*^fl/fl^ and CD4-Cre *Panx1*^fl/fl^ mice, with introduction of CD45.1 and CD45.2 congenic markers for identification. Animals were maintained under specific-pathogen-free conditions at Mayo Clinic Arizona and at the University of Minnesota. In all experiments, mice were randomly assigned to experimental groups. All experimental procedures were approved by the institutional animal care and use committee at Mayo Clinic Arizona (IACUC A00005542-20) and at the University of Minnesota (IACUC 1709-35136A).

### Viral strains

LCMV (Armstrong strain) was maintained at −80°C until infection and diluted to 2x10^6^ PFU/ml in PBS. Influenza (PR8 strain) was maintained at −80°C and diluted to 7x10^4^ PFU/ml in PBS at the time of infection studies.

### Infection studies

WT (CD4-Cre, ERT2-Cre-LSL-YFP), CD4-Cre *Panx1*^fl/fl^, ERT2-Cre-LSL-YFP-*Panx1*^fl/fl^, and CD4-Cre *P2rx7*^fl/fl^ P14 cells were adoptively transferred into naive wild-type mice, which were infected with LCMV-Armstrong (2 × 10^5^ PFU, intraperitoneally (i.p.)). Sometimes, CD4-Cre or CD4-Cre *Panx1*^fl/fl^ mice were infected with LCMV Armstrong. In other experiments, CD4-Cre or CD4-Cre *Panx1*^fl/fl^ mice were infected with Influenza-PR8 (1400 PFU, intranasally (i.n.)). In some experiments, LCMV-infected mice were treated with Trovafloxacin (4.2 mg x kg^-1^ mouse, Sigma-Aldrich) or Sodium Lactate (1.68 g x kg^-1^ mouse, Sigma-Aldrich)^69^ between days 1-7 after infection.

### Tumor inoculations

WT or CMV-Cre *Panx1^f^*^l/fl^ mice were injected subcutaneously with 1.5 x 10^5^ B16.F10 or 3 x 10^5^ B16.F10.gp33 cells in the right flank. In some experiments, after tumors became palpable (∼7 days), tumor-bearing mice received 5 x 10^5^ activated CD4-Cre or CD4-Cre *Panx1*^fl/fl^ P14 cells (or 2.5 x 10^5^ of a 1:1 mix of these two P14 populations for co-transfer experiments). Tumor growth was monitored by measuring height and width with calipers until mice reached an endpoint criterion of 144 mm^2^ or ulceration. When indicated in the figures, tumor-bearing mice were sacrificed for flow cytometry analysis.

### Bone marrow chimeras

CD45.1/2^+^ WT mice were reconstituted with a 1:1 mixture of CD4-Cre (CD45.1^+^) and CD4-Cre *Panx1*^fl/fl^ (Panx1-KO, CD45.2^+^) following lethal irradiation (2 x 450 Rad). Two months following irradiation, mice were infected with LCMV-Armstrong.

### GVHD mouse model

GVHD mouse experiments were done as described previously (PMID: 11850710). BALB/c mice were lethally irradiated (2 x 450 Rad) and reconstituted with either bone marrow cells from BALB/c or C57BL/6 mice. In the mice transferred with C57BL/6 BM cells (1 x 10^7^ cells), spleen CD8^+^ T cells from either CD4-Cre or CD4-Cre *Panx1*^fl/fl^ mice were transferred together (2 x 10^6^ cells). Weight loss was monitored in recipient BALB/c mice for 7 days.

### Administration of tamoxifen in mice

Tamoxifen (Sigma-Aldrich, diluted in Sunflower Oil) was administered to mice i.p. at a daily dose of 1 mg ^37^. Two different treatment periods were used, in which Tamoxifen was given between days 1-4 or between days 28-35 after LCMV-Armstrong infection.

### Primary cell cultures

Total CD8^+^ T cells or P14 cells were obtained from 6- to 8-week-old mice with a C57BL/6 background. Cells were cultured in complete RPMI media: RPMI 1640 (Corning) supplemented with 10% Fetal Bovine Serum (Atlanta Biologicals), 100 U/ml penicillin/streptomycin (Thermo Fisher Scientific) and 2 mM L-glutamine (Corning). All cells were cultured at 37°C in a humidified atmosphere containing 5% CO_2_.

### *In vitro* activation of CD8^+^ T cells

P14 and/or polyclonal WT (CD4-Cre), WT Nur77-GFP, P2RX7-KO (CD4-Cre *P2rx7*^fl/fl^) or Panx1-KO (CD4-Cre *Panx1*^fl/fl^) CD8^+^ T cells were isolated from naïve mice with the Mouse CD8^+^ T cell isolation kit (Miltenyi Biotec). Cells were stimulated for 3h, 24h, 48h or 72h with plate-bound αCD3 (10 μg/ml), plate-bound αCD28 (20 μg/ml) and soluble IL-2 (10 ng/ml). When indicated, ATP (50 μM, Sigma-Aldrich), Trovafloxacin (Sigma-Aldrich, 10 μM), A-438079 (Tocris, 10 μM), 5-BDBD (Sigma-Aldrich, 10 μM), Sodium Lactate (1 mM, Sigma-Aldrich), VB-124 (10 μM, Tocris), Oligomycin (1.39 nM, Agilent), BSA-conjugated Oleic Acid (100 μM, ThermoFisher), Inostamycin (0.2 μg/ml, Cayman Chemical), Glucose (Sigma-Aldrich, 1 mM) or SR13800 (10 μM, Cayman Chemical) were added to the cultures at the beginning of activation. In screening experiments, multiple other compounds were added to the cultures, or in alternative their respective vehicles. In some experiments, after 72h of activation, CD8^+^ T cells were incubated for an additional 72h with either recombinant IL-2 (10 ng/ml) or IL-15 (10 ng/ml). In some experiments, CD4-Cre or CD4-Cre *Panx1*^fl/fl^ CD8^+^ T cells were activated *in vitro* for 3h with PMA and Ionomycin (Cell Stimulation Cocktail, ThermoFisher).

### Western Blot

The indicated CD8^+^ T cells were lysed in RIPA buffer supplemented with 1 mM PMSF and protease/phosphatase inhibitors. Cell lysates were then sonicated, and protein concentration measured using colorimetric BCA assay. Aliquots of 25 μg of protein were run on 4-12% gradient agarose gels and transferred to nitrocellulose membranes using a Trans-Blot Turbo system. Membranes with transferred proteins were blocked for 30 minutes at room temperature with TBS, then stained with primary rabbit anti-Pannexin-1 (Cell Signaling Technologies) at 1/1000 dilution or mouse anti-Tubulin (Millipore) at 1/3000 dilution, at 4°C, overnight with rotation. After washing, membranes were stained with secondary anti-rabbit or anti-mouse antibodies (1/15000 dilution, 1h at room temperature). After washing, membranes were imaged using a LICOR Odyseey DLx system.

### Flow cytometry

Lymphocytes were isolated from tissues including thymus, spleen, inguinal lymph nodes, cervical lymph nodes, blood, small intestine epithelium (SI IEL), salivary glands (SG), tumors, lungs and kidneys as previously described ^32,37,60^. In summary, organs were removed and cut in small pieces into Erlenmeyer flasks containing 30 mL of 0.5 mg/ml Collagenase type I solution (tumor, kidney, and SG), 0.5 mg/ml Collagenase type IV (lung) or 0.15 mg/ml Dithioerythritol (SI IEL). Following this period, lymphocytes were isolated by 44/67% Percoll gradient isolation. During isolation of lymphocytes from non-lymphoid tissues, in all experiments, 50 μg of Treg-Protector (anti-ARTC2.2) nanobodies (BioLegend) were injected i.v. 30 minutes prior to mouse sacrifice ^35^. Direct *ex vivo* staining and intracellular staining were performed as described ^32^. To identify LCMV-specific or Flu-specific CD8^+^ T cells, tetramers were obtained from the Yerkes NIH Tetramer Core: D^b^-gp33 and D^b^-NP-flu tetramers conjugated with APC-or PE-Streptavidin were used. For detection of vascular-associated lymphocytes in non-lymphoid organs, *in vivo* i.v. injection of PerCP-Cy5.5-conjugated CD8α antibody was performed ^81^. Among LCMV-or Flu-specific CD8^+^ T cells, the following markers were used to distinguish these respective populations: naïve (CD44^-^CD62L^+^), Tcm (CD44^+^CD62L^+^), Tem (CD44^+^CD62L^-^CD127^+^), LLEC (CD44^+^CD62L^-^CD127^-^KLRG1^hi^), Trm (i.v.CD8α^-^CD69^+/−^CD103^hi/int/lo^), MPs (CD127^+^KLRG1^-)^, TEs (CD127^-^KLRG1^+^). Adoptively transferred P14 cells and LCMV-specific CD8^+^ T cells in mixed bone marrow chimeras were identified using a combination of CD45.1 and CD45.2 monoclonal antibodies. In some experiments, To-Pro 3 (Sigma-Aldrich) was added right before flow cytometry ^22^. From *in vitro*-activated CD8^+^ T cells, we used antibodies for CD44, CD69, CD25; in some experiments, CD8^+^ T cells were stained with Cell Trace Violet (Life Technologies) according to manufacturer’s instructions, for tracking of proliferation cycles. In all flow cytometry experiments, Live/Dead Near-IR was used to distinguish between live and dead cells. Intracellular detection of Ki-67 and IFNγ was done as described before ^32^. For mitochondrial mass and membrane potential measurements, cells were incubated with Mito Green (Thermo Fisher Scientific) and Mitotracker Deep Red (Thermo Fisher Scientific) or TMRE (Cell Signaling Technology) simultaneously for 15 min at 37°C prior to staining. For intracellular detection of phosphorylated Acetyl-CoA Carboxylase (pACC; Cell Signaling Technology) or phosphorylated ERK1/2 (pERK1/2; Biolegend), cells were stained with surface markers then fixed with Paraformaldehyde 1%, permeabilized with 90% Methanol and stained with pACC or pERK1/2-PE for 20 min at room temperature. A secondary antibody (Alexa Fluor 647 anti-rabbit IgG) was used for detection of pACC. Alexa Fluor 488-conjugated Cholera Toxin B was used for detection of lipid rafts and stained as described in ^49^. Nur77 was detected as GFP signal from Nur77-GFP cells via flow cytometry. Apoptosis (cleaved Caspase 3/7) measurements were done using the FMK-FLICA-Caspase 3/7 Kit (+ PI staining; Thermo Fisher Scientific) following the manufacturer instructions. Flow cytometric analyses were performed on FACS Symphony or LSR Fortessa (BD Biosciences) and data was analyzed using FlowJo software (Treestar).

### Cell sorting

Cell sorting was performed on a FACS Aria III device (BD Biosciences). RNA-seq analysis experiments were performed on WT (CD4-Cre) and Panx1-KO (CD4-Cre *Panx1*^fl/fl^) TE (KLRG1^+^CD127^-^) and MP (KLRG1^-^CD127^+^) P14 cells sorted from mice 7 days post-LCMV infection. The population purity after cell sorting was > 98% in all experiments.

### Intracellular and extracellular ATP measurements

Intracellular ATP concentrations were measured by using the ATP determination kit (Life Technologies). To assess the levels of extracellular ATP, WT CD8^+^ T cells were activated *in vitro* as described above, and further incubated with IL-2 for 24h; IL-2 incubation cultures were performed in Transwells (Corning). After 24 h of IL-2 incubation, the transwells were transferred to new wells and the plates were centrifuged at 50 rpm for 1 min. The supernatant below the transwells was quantified for ATP concentration as described above. In all transwells, a combination of the plasma membrane ATPase inhibitor Ebselen (30 μM; Cayman Chemical) and the ecto-ATPase inhibitor ARL 67156 trisodium salt hydrate (100 μM; Sigma-Aldrich) was added 1 h before evaluation, to inhibit ATPase activity ^32^. In some samples, Trovafloxacin (10 μM) was added 24 h before assessment.

### CRISPR-Cas9 experiments

Cas9/RNP nucleofection of P14 cells was performed as previously described^53^. Briefly, WT (CD4-Cre) and CD4-Cre *P2rx7*^fl/fl^ P14 cells were isolated and activated as described above. After 24h of activation, single guide RNAs (sgRNA) for either *Cd4* (sg*Cd4*), *Panx1* (sg*Panx1*) or *P2rx4* (sg*P2rx4*) and Cas9 protein were pre-complexed at room temperature for at least 10 mins. Following this, 1-10 million of pre-activated P14 cells were resuspended in 20 μl of P4 Nucleofection Buffer (Lonza). This cell suspension was then mixed with sgRNA/Cas9 and then incubated at room temperature for 2 mins. The P14 cell/Cas9/RNP mixes were transferred to Nucleofection cuvette strips (4D-Nucleofector X kit S; Lonza). Cells were electroporated using a 4D nucleofector (4D-Nucleofector Core Unit; Lonza). After nucleofection, prewarmed complete RPMI was used to transfer transfected P14 cells in 96-well plates. After 2h, P14 cells were cultured in 24-well plates in complete RPMI for 48 h, before transfer into recipient mice.

### RNA-seq and bioinformatics analyses

WT (CD4-Cre) or CD4-Cre *Panx1*^fl/fl^ P14 cells were *in vitro* activated for 72h (“effector-like”) or activated and subsequently treated with recombinant IL-15 (“memory-like”). In another experiment, CD4-Cre or CD4-Cre *Panx1*^fl/fl^ TE or MP cells were sorted from mice infected with LCMV-Arm (7 days p.i.). These cells were homogenized using QIAshredder columns (Qiagen), and RNA was extracted using the RNeasy Plus Mini Kit (Qiagen) following the manufacturer’s instructions. Library preparation and RNA-seq (BGI-Seq platform, PE 100-bp paired-end read length) was done by BGI Americas. RNA-seq reads were mapped, and raw count matrix was generated. DEG analysis was done using DESeq2, and genes with greater than twofold changes and false discovery rate <0.05 were considered for gene cluster analysis. BGI provided customized analysis of DEGs, principal component analysis plots, and Pathway analyses between groups. Some heatmaps were generated using the Morpheus software (Broad Institute; https://software.broadinstitute.org/morpheus/). Some RNA-seq expression value counts were collected from the public database ImmGen (immgen.org), and from previous RNA-seq analysis performed by our group ^32,34^. These RNA-seq analyses can be accessed at GEO Omnibus with the accession numbers GSE217292 and GSE218643. Analysis of single-cell RNA-seq from spleen P14 cells from days 3 or 7 after LCMV-Arm infection^82^ (GSE131847) was done using R Studio.

### Extracellular flux analyses

In the indicated naïve, *in vitro*-activated or *ex vivo*-sorted CD8^+^ T cell populations, OCR and ECAR values were measured using a 96-well XF Extracellular flux analyzer, following manufacturer instructions (Agilent). Spare respiratory capacity (SRC) was defined as the subtraction between the maximum OCR values and the baseline OCR values ^32,37^.

### Metabolomics analyses

For untargeted metabolomics, the indicated *in vitro*-activated CD8^+^ T cell populations were isolated. We separated cell pellets and supernatants and sent them to the Mayo Clinic Metabolomics Core Laboratory for Liquid Chromatography-Mass Spectrometry (LC-MS) metabolome assessments. Primary analyses and comparisons of differentially expressed metabolites (DEMs) were done by the Metabolomics Core. Pathway enrichment analyses of DEMs were done using the MetaboAnalyst tool. Heatmaps of the top-100 DEMs for each comparison were generated using the Morpheus software. For targeted metabolomics analyses, *in vitro*-activated CD8^+^ T cells will be cultured for the final 12h of activation with ^13^C-Glucose (Cambridge Isotopes) or ^13^C-Sodium Lactate (Cambridge Isotopes). We separated cell pellets and sent them to the Mayo Clinic Metabolomics Core Laboratory for metabolome assessments of the TCA cycle.

### L-lactate detection assays

When indicated, the supernatants from *in vitro*-activated CD8^+^ T cell populations or serum from LCMV-infected mice were collected, and the concentrations of extracellular L-lactate were measured using the L-Lactate Assay Kit (Sigma-Aldrich), following the manufacturer’s instructions.

### Quantification and statistical analyses

Details on statistics used can be found in figure legends. Data were subjected to the Kolmogorov-Smirnov test to assess normality of samples. Statistical differences were calculated by using unpaired two-tailed Student’s t-test, one-way ANOVA with Tukey post-test, two-way ANOVA with Bonferroni’s post-test, or Kaplan-Meier survival curve analysis. All experiments were analyzed using Prism 10 (GraphPad Software). Graphical data was shown as mean values with error bars indicating the SD. P values of < 0.05 (*), < 0.01 (*), < 0.001 (***) or < 0.0001 (****) indicated significant differences between groups.

## Supplementary Materials

This article includes Figs.S1 to S7.

