## Supplementary Figures for "The ATP-exporting channel Pannexin-1 promotes CD8^+^ T cell effector and memory responses"

**Supplemental information titles and legends**

**Figure S1. Gating strategy for flow cytometry, related to Figures 1-7.** (**a**) Gating for identification of $CD8^+$ T cells. (**b**) Gating for identification of $gp33^+$ $CD8^+$ T cells. (**c**) Gating for identification of P14 cells in recipient mice. (**d**) Gating for identification of TE and MP cells. (**e**) Gating for identification of circulating (left) and tissue-resident (right) memory $CD8^+$ T cell subsets. (**f**) Gating for the ERT2-Cre experiments.

**Figure S2. Functional Panx1 hemichannels are expressed by $CD8^+$ T cells and are dispensable for the thymic development of $CD8^+$ T cells, related to Figure 1.** (**a**) Panx1 mRNA expression (data from *Immgen.org*) in the indicated subsets of P14 cells. (**b**) $CD8^+$ T cells were *in vitro* activated (αCD3/αCD28/IL-2); after 48h, vehicle or Trovafloxacin was added to the activation cultures. The ATP concentration of the supernatants of the pericellular space (eATP) and the intracellular lysate of $CD8^+$ T cells (iATP) were quantified. (**c**) WT and global Panx1-KO (CMV-Cre *Panx1*$^{fl/fl}$) mice were infected with LCMV-Arm; at day 7 post-infection, spleens were collected and incubated with To-Pro 3 dye. Histograms showing the internalization of To-Pro 3 in WT versus Panx1-KO $gp33^+$ $CD8^+$ T cells are shown. As positive control of To-Pro 3 internalization, Live/Dead$^+$ $CD8^+$ T cells from WT mice is represented. (**d-e**) CD4-Cre and CD4-Cre *Panx1*$^{fl/fl}$ mice were harvested for thymus and spleen cell analyses. (**d**) Flow cytometry plots showing expression of CD4 and CD8a in thymocytes (left); in the right, average percentages of CD4/CD8 double-negative (DN), double-positive (DP), $CD4^+$ (CD4 SP) and $CD8^+$ (CD8 SP) thymocytes. (**e**) Average numbers of $CD4^+$ and $CD8^+$ T cells in the spleen. (**f**) Single-cell RNA-seq (scRNAseq) analysis of effector P14 cells showing a representative heatmap with genes defining the UMAP clusters depicted in Figure 1 (left); in the right, the expression of Klrg1 and Sell mRNA are shown for each cluster. (**g**) Full blots for Pannexin-1 (left) and Tubulin (right) expression, related to Figure 1c. (**b-e, g**) Data from 2-3 independent experiments, n=3-9 per experimental group. ns – not significant (p>0.05), ****p<0.0001, Unpaired t-test (**b**), One-way ANOVA with Tukey's post-test (**d-e**).

**Figure S3. Panx1 pharmacological inhibition impacts CD8$^+$ T cell effector immune responses *in vivo*, related to Figure 1.** (**a**) C57BL/6 mice (adoptively transferred with P14 cells and infected with LCMV-Arm) were treated between days 1-7 post-infection (p.i.) with Vehicle or Trovafloxacin. Numbers of P14 cells per spleen, at day 7 p.i., are shown. (**b**) *Panx1* mRNA expression (data from *Immgen.org*) in the indicated cell subsets. (**a**) Data from two independent experiments, n=3-5 per experimental group, ** p<0.01, Unpaired t-test.

**Figure S4. Panx1 promotes CD8$^+$ T cell activation and proliferation in a partially autocrine way, related to Figures 1 and 2.** (**a-b**) P14 cells (CD45.2$^+$) were *in vitro* activated and transduced via electroporation with a mix of Cas9 and either single-guide RNAs for *Cd4* (sg*Cd4*) or *Panx1* (sg*Panx1*). After a 48h rest period, cells were adoptively transferred into CD45.1$^+$ recipient mice, which were infected with LCMV-Arm. (**a**) Representative flow cytometry plots showing expression of KLRG1 and CD127 in spleen sg*Cd4* and sg*Panx1*-P14 cells, at day 7 post-infection. (**b**) Percentages of TE and MP WT P14 cells (right) at day 7 post-infection. (**c-d**) In other experiments, CD4-Cre or CD4-Cre *Panx1*$^{fl/fl}$ mice were infected with Influenza virus (PR8 strain). At 7 days p.i., the spleens and lung parenchyma (lung i.v.$^-$) were assessed. (**c**) Average numbers of NP-tetramer$^+$ CD8$^+$ TE and MP cells per spleen. (**d**) Average numbers of NP-tetramer$^+$ CD8$^+$ T cells per lung. (**e-f**) CD4-Cre (WT; CD45.1$^+$) and CD4-Cre *Panx1*$^{fl/fl}$ (KO; CD45.2$^+$) CD8$^+$ T cells were activated alone or co-cultured (1:1) and activated *in vitro*. (**e**) Experimental design. (**f**) Average CD44 gMFI values (left) and percentages of CTV divided cells (right) at 72h post-activation. (**a-f**) Data from 2-3 independent experiments, n=3-18 per experimental group. ns – not significant (p>0.05), * p<0.05, ** p<0.01, **** p<0.0001, One-way ANOVA with Tukey's post-test (**b-c, f**), Unpaired t-test (**d**).

**Figure S5. Panx1 effects on the extracellular and intracellular metabolism of CD8$^+$ T cells, related to Figures 2-4.** (**a**) WT (CD4-Cre) or Panx1-KO (CD4-Cre *Panx1*$^{fl/fl}$) P14 cells (CD45.2$^+$) were transferred into LCMV-infected WT CD45.1$^+$ mice, which were analyzed at day 3 post-infection. Flow cytometry plots showing Mito Green and TMRE expression (left) and average values of Mito Green gMFI, TMRE gMFI and percentages of depolarized mitochondria are shown. (**b**) CD4-Cre or CD4-Cre *Panx1*$^{fl/fl}$ effector-like and memory-like CD8$^+$ T cell supernatants were harvested, and the levels of eATP were measured. (**c**) CD4-Cre or CD4-Cre *Panx1*$^{fl/fl}$ CD8$^+$ T cells were activated *in vitro* (anti-CD3/CD28 + IL-2) for 3h, in the presence of eATP +/- 5-BDBD (P2RX4i); average percentages of CD69$^+$ cells (left), CD25$^+$ cells (middle) and CD44$^+$ cells (right) are shown. (**d**) P14 cells (CD45.2$^+$) were *in vitro* activated and transduced via electroporation with a mix of Cas9 and either single-guide RNAs for *Cd4* (sg*Cd4*) or *P2rx4* (sg*P2rx4*). After a 48h rest period, cells were adoptively transferred into CD45.1$^+$ recipient mice, which were infected with LCMV-Arm. The average numbers of spleen P14 cells per mouse are shown. (**e-g**) WT (CD4-Cre) or Panx1-KO (CD4-Cre *Panx1*$^{fl/fl}$) effector-like and memory-like CD8$^+$ T cell cultures were harvested, and intracellular lysates and supernatants were submitted for untargeted metabolomics (GC-MS) analysis. (**e**) Experimental design. (**f**) PCA plots comparing WT and Panx1-KO CD8$^+$ T cell samples. (**g**) Heatmaps showing the top 100 differentially expressed metabolites (DEMs) with increased representation in the supernatants of effector-like (left) or memory-like (right) WT cells, in comparison to Panx1-KO cells. (**h**) mRNA counts per million of *Slc16a1* (MCT-1-encoding gene) and *Slc16a3* (MCT-4-encoding gene) genes in CD4-Cre (blue) and CD4-Cre *Panx1*$^{fl/fl}$ (red) effector-like cells. (**i**) In some experiments, the top DEMs depicted in Fig. S5f (effector-like; left) were added exogenously in WT (CD4-Cre) versus Panx1-KO (CD4-Cre *Panx1*$^{fl/fl}$) CD8$^+$ T cell *in vitro* activation cultures. The percentage of Panx1-KO activation rescue (inferred as percentage of increase in CD69 expression compared to WT cells) is depicted for these metabolites. (**j**) Percentages, within total intracellular PIs, of saturated (one or less double bonds) and unsaturated (2 or more double bonds) PIs in WT versus Panx1-KO effector-like CD8$^+$ T cells. Data from the GC-MS experiments described above. (**k**) CD4-Cre and CD4-Cre Panx1$^{fl/fl}$ CD8$^+$ T cells were *in vitro* activated in the presence or not of sodium lactate and/or SR13800 (MCT1i); the average gMFI values of Cholera Toxin B per group are shown. (**l**) Intracellular levels (a.u.) of oleic acid in WT versus Panx1-KO effector-like CD8$^+$ T cells (from the GC-MS experiments). (**m**) CD4-Cre and CD4-Cre *Panx1*$^{fl/fl}$ CD8$^+$ T cells were activated in the presence or not of sodium lactate and/or Inostamycin (CDIPTi); the average gMFI values of Cholera Toxin B per group are shown. (**a-d, i, k, m**) Data from 2-3 independent experiments, n=3-6 per experimental group. (**e-h, j, l**) Data from 2-3 biological replicates per experimental group (n=3 per replicate). ns – not significant (p>0.05), * p<0.05, ** p<0.01, ***p<0.001, **** p<0.0001, Unpaired t-test (**a, d, j, l**), One-way ANOVA with Tukey's post-test (**b-c, h-i, k, m**).

**Figure S6. Panx1-KO *in vivo* effector CD8$^+$ T cell expansion is rescued by exogenous sodium lactate, related to Figure 3.** (**a-e**) WT (CD4-Cre) or Panx1-KO (CD4-Cre *Panx1*$^{fl/fl}$) P14 cells (CD45.2$^+$) were transferred into WT CD45.1$^+$ mice, which were infected with LCMV-Arm. Some mice were treated with sodium lactate between days 1-3 post-infection. Spleen P14 cells were analyzed at day 7 post-infection. (**a**) Experimental design. (**b**) Average concentrations of lactate in the serum of mice at day 4 post-infection. (**c**) Representative flow cytometry plots showing P14 cell percentages (CD45.2$^+$) in the indicated groups. (**d**) Average numbers of total spleen P14 cells at day 7 post-infection. (**e**) Average percentages of TE, MP and DN P14 cells per spleen are shown. (**b, d-e**) Data from two independent experiments, n=3-5 per time point. (**b, d-e**) ns – not significant, * p<0.05, **** p<0.0001, Unpaired t-test (**b**), One-way ANOVA with Tukey's post-test (**d-e**).

**Figure S7. Effects of Panx1 deletion on memory-phenoype CD8$^+$ T cells, related to Figures 5-6.** (**a-c**) Recipient C57BL/6 mice (CD45.1$^+$) were adoptively transferred with a 1:1 mix of P14 ERT2-Cre LSL-YFP *Panx1*$^{+/+}$ (WT; CD45.1/2$^+$) and P14 ERT2-Cre LSL-YFP *Panx1*$^{fl/fl}$

(CD45.2$^+$) cells, then infected with LCMV-Arm. Infected mice were treated with vehicle or tamoxifen (Tx) between days 1-4 post-infection (Tx1-4). (**a**) *Panx1*$^{fl/fl}$/*Panx1*$^{+/+}$ P14 cell ratios in the blood over time post-infection, in mice treated with vehicle or Tx1-4. Treatment period is indicated by the blue shaded box. (**b**) *Panx1*$^{fl/fl}$/*Panx1*$^{+/+}$ ratios of spleen $T_{CM}$, $T_{EM}$ and LLEC P14 cells from mice treated with vehicle or Tx1-4, at day 63 post-infection. (**c**) *Panx1*$^{fl/fl}$/*Panx1*$^{+/+}$ P14 cell ratios in SG and SI IEL at day 63 post-infection, in mice treated with vehicle or Tx1-4. (**d**) ECAR kinetics for sorted CD4-Cre (blue) and CD4-Cre *Panx1*$^{fl/fl}$ (red) P14 MP cells. (**e**) ECAR kinetics for CD4-Cre (blue) and CD4-Cre *Panx1*$^{fl/fl}$ (red) memory-like CD8$^+$ T cells. (**a-e**) Data from 2-3 independent experiments, n=5-12 per experimental group. ns – not significant ($p>0.05$), *$p<0.05$, **$p<0.01$, ***$p<0.001$, ****$p<0.0001$, Two-way ANOVA with Bonferroni's post-test (**a**), One-way ANOVA with Tukey's post-test (**b-c**).

### Figure S1

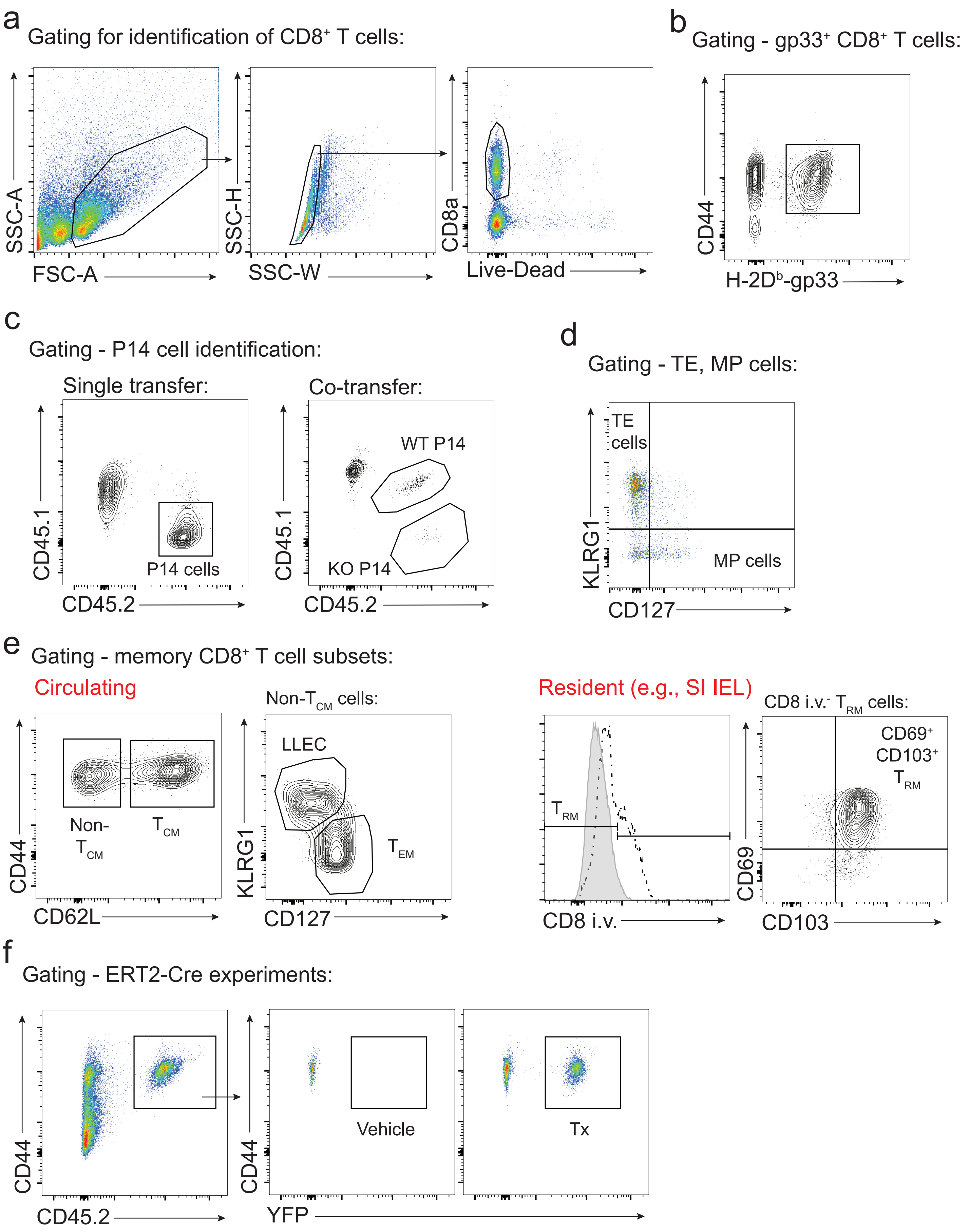
a
Gating for identification of CD8+ T cells:
SSC-A
FSC-A
SSC-H
SSC-W
CD8a
Live-Dead
b
Gating - gp33+ CD8+ T cells:
CD44
H-2Db-gp33
c
Gating - P14 cell identification:
Single transfer:
CD45.1
CD45.2
P14 cells
Co-transfer:
WT P14
KO P14
CD45.1
CD45.2
d
Gating - TE, MP cells:
TE cells
MP cells
KLRG1
CD127
e
Gating - memory CD8+ T cell subsets:
Circulating
Non-TCM
TCM
CD44
CD62L
Non-TCM cells:
LLEC
TEM
KLRG1
CD127
Resident (e.g., SI IEL)
TRM
CD8 i.v.
CD8 i.v.- TRM cells:
CD69+ CD103+ TRM
CD69
CD103
f
Gating - ERT2-Cre experiments:
CD44
CD45.2
Vehicle
Tx
CD44
YFP

### Figure S2

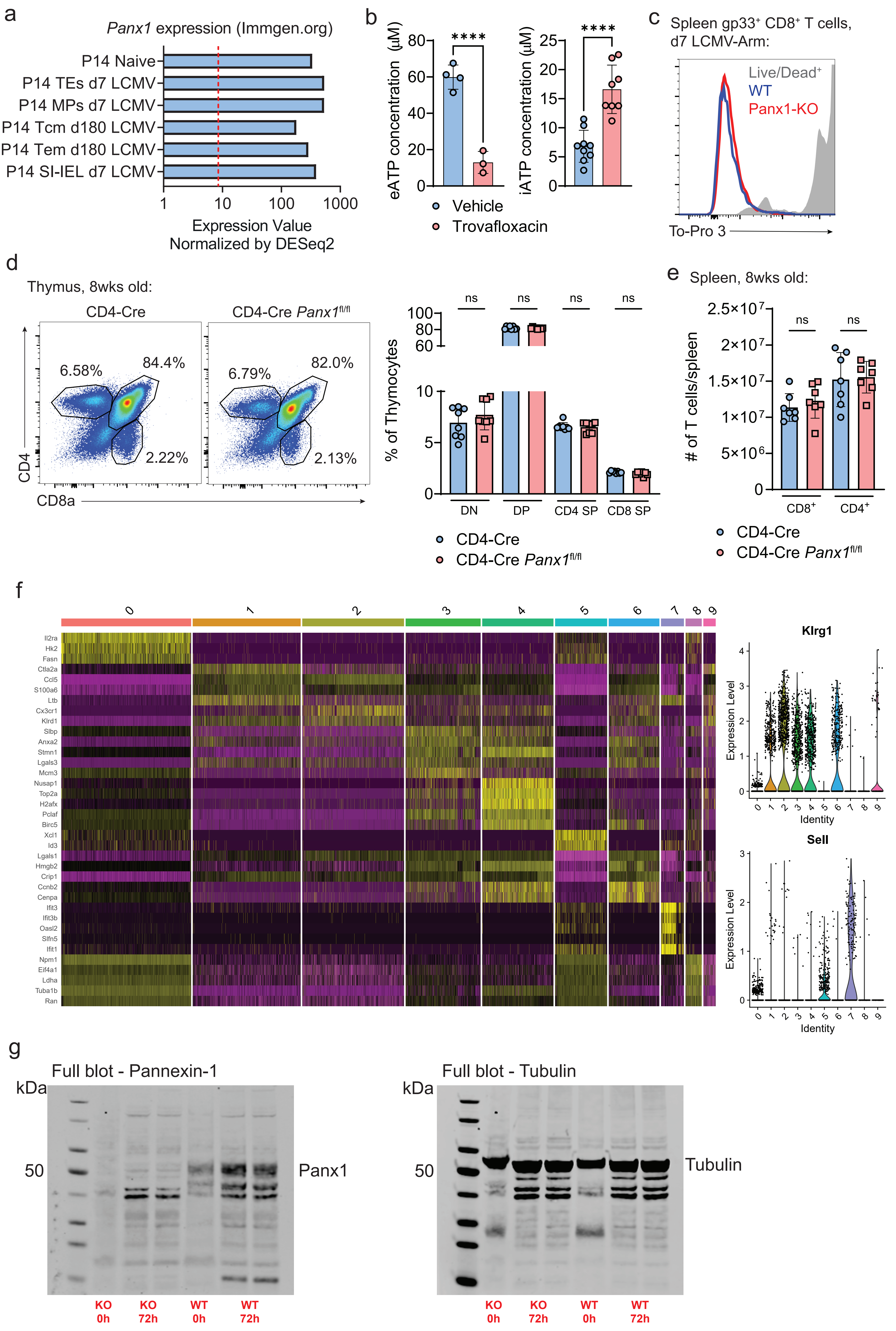
a
Panx1 expression (Immgen.org)
P14 Naive
P14 TEs d7 LCMV
P14 MPs d7 LCMV
P14 Tcm d180 LCMV
P14 Tem d180 LCMV
P14 SI-IEL d7 LCMV
1
10
100
1000
Expression Value
Normalized by DESeq2
b
eATP concentration (μM)
iATP concentration (μM)
****
****
Vehicle
Trovafloxacin
c
Spleen gp33+ CD8+ T cells, d7 LCMV-Arm:
Live/Dead+
WT
Panx1-KO
To-Pro 3
d
Thymus, 8wks old:
CD4-Cre
CD4-Cre Panx1fl/fl
6.58%
84.4%
2.22%
6.79%
82.0%
2.13%
CD4
CD8a
% of Thymocytes
ns
DN
DP
CD4 SP
CD8 SP
CD4-Cre
CD4-Cre Panx1fl/fl
e
Spleen, 8wks old:
### of T cells/spleen
2.5×10^7
2×10^7
1.5×10^7
1×10^7
5×10^6
ns
CD8+
CD4+
CD4-Cre
CD4-Cre Panx1fl/fl
f
Il2ra
Hk2
Fasn
Ctla2a
Ccl5
S100a6
Ltb
Cx3cr1
Klrd1
Slbp
Anxa2
Stmn1
Lgals3
Mcm3
Nusap1
Top2a
H2afx
Pclaf
Birc5
Xcl1
Id3
Lgals1
Hmgb2
Crip1
Ccnb2
Cenpa
Ifit3
Ifit3b
Oasl2
Slfn5
Ifit1
Npm1
Eif4a1
Ldha
Tuba1b
Ran
Klrg1
Sell
Expression Level
Identity
g
Full blot - Pannexin-1
Full blot - Tubulin
kDa
50
Panx1
Tubulin
KO 0h
KO 72h
WT 0h
WT 72h

### Figure S3

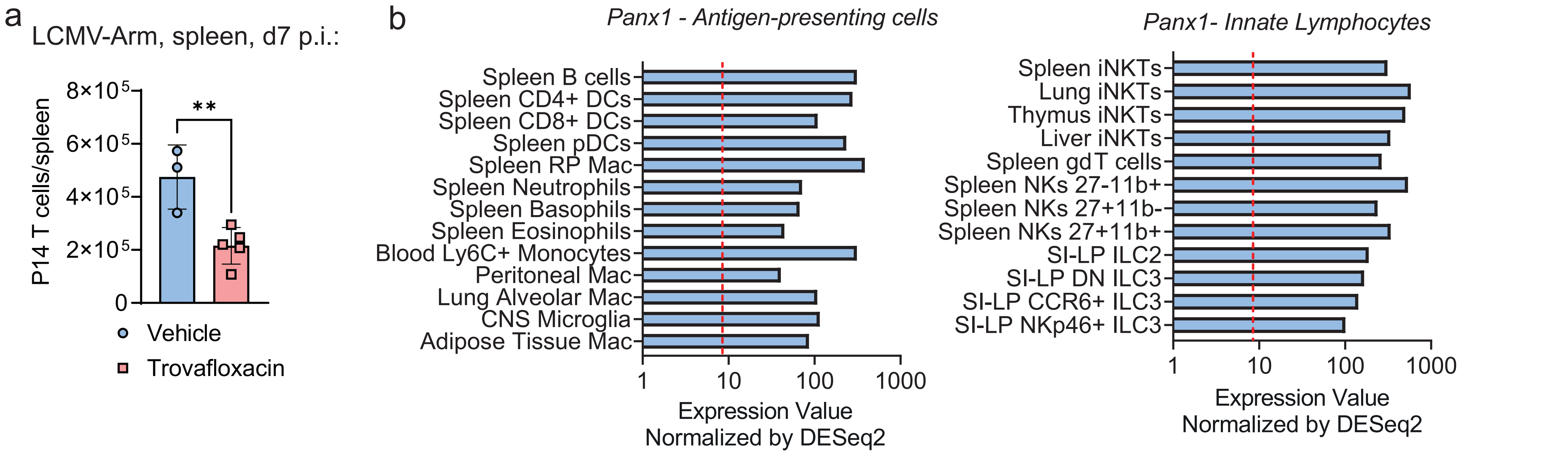
a
LCMV-Arm, spleen, d7 p.i.:
P14 T cells/spleen
8×10^5
6×10^5
4×10^5
2×10^5
0
**
Vehicle
Trovafloxacin
b
Panx1 - Antigen-presenting cells
Spleen B cells
Spleen CD4+ DCs
Spleen CD8+ DCs
Spleen pDCs
Spleen RP Mac
Spleen Neutrophils
Spleen Basophils
Spleen Eosinophils
Blood Ly6C+ Monocytes
Peritoneal Mac
Lung Alveolar Mac
CNS Microglia
Adipose Tissue Mac
1
10
100
1000
Expression Value
Normalized by DESeq2
Panx1- Innate Lymphocytes
Spleen iNKTs
Lung iNKTs
Thymus iNKTs
Liver iNKTs
Spleen gd T cells
Spleen NKs 27-11b+
Spleen NKs 27+11b-
Spleen NKs 27+11b+
SI-LP ILC2
SI-LP DN ILC3
SI-LP CCR6+ ILC3
SI-LP NKp46+ ILC3
1
10
100
1000
Expression Value
Normalized by DESeq2

### Figure S4

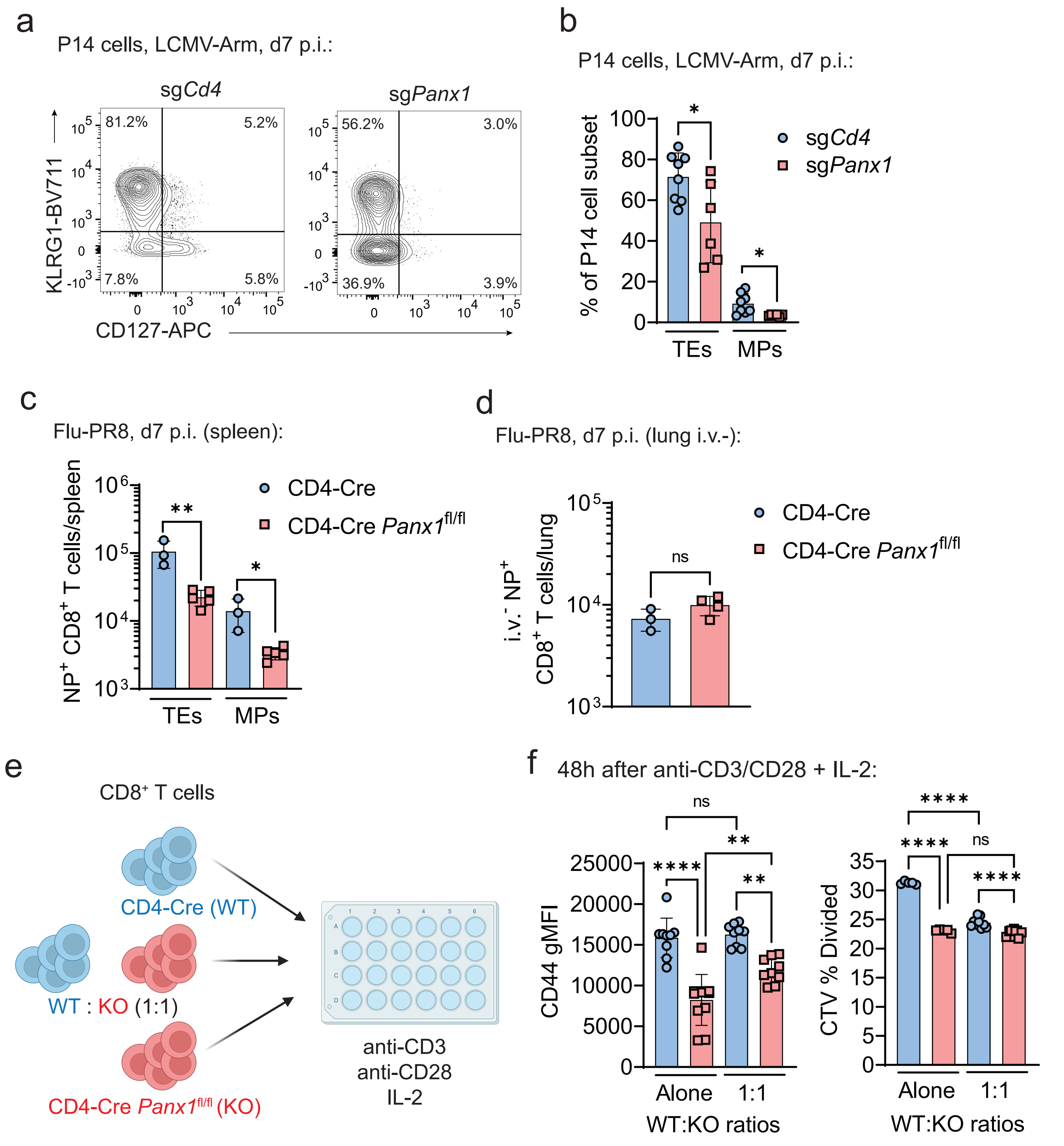
a
P14 cells, LCMV-Arm, d7 p.i.:
sgCd4
sgPanx1
81.2%
5.2%
7.8%
5.8%
56.2%
3.0%
36.9%
3.9%
KLRG1-BV711
CD127-APC
b
P14 cells, LCMV-Arm, d7 p.i.:
% of P14 cell subset
sgCd4
sgPanx1
TEs
MPs
c
Flu-PR8, d7 p.i. (spleen):
NP+ CD8+ T cells/spleen
CD4-Cre
CD4-Cre Panx1fl/fl
TEs
MPs
d
Flu-PR8, d7 p.i. (lung i.v.-):
i.v.- NP+ CD8+ T cells/lung
ns
CD4-Cre
CD4-Cre Panx1fl/fl
e
CD8+ T cells
CD4-Cre (WT)
WT : KO (1:1)
CD4-Cre Panx1fl/fl (KO)
anti-CD3
anti-CD28
IL-2
f
48h after anti-CD3/CD28 + IL-2:
CD44 gMFI
CTV % Divided
ns
Alone
1:1
WT:KO ratios

### Figure S5

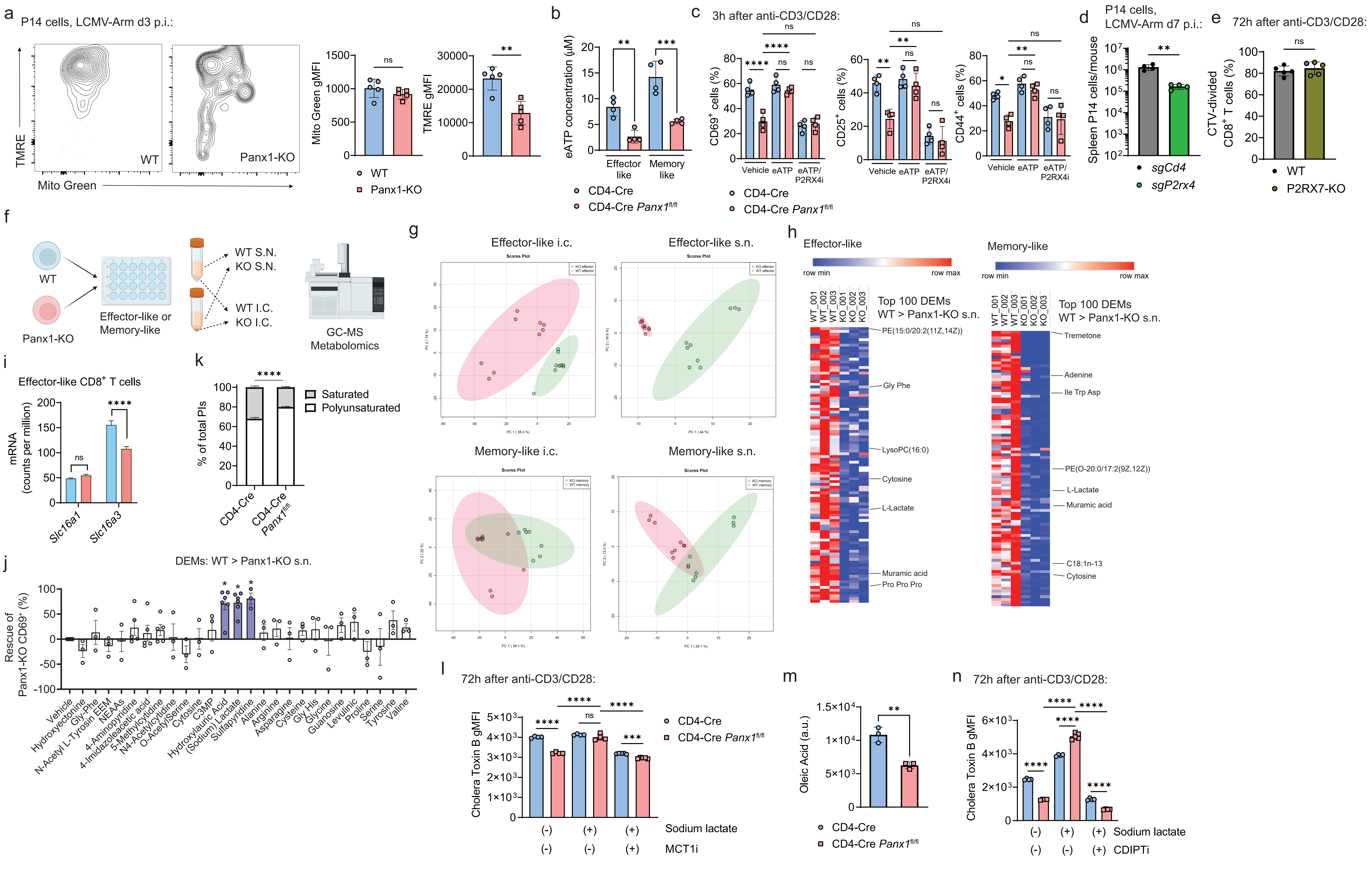

a
P14 cells, LCMV-Arm d3 p.i.:
TMRE
Mito Green
WT
Panx1-KO
Mito Green gMFI
TMRE gMFI
ns
**
WT
Panx1-KO
b
eATP concentration (μM)
**
***
Effector like
Memory like
CD4-Cre
CD4-Cre Panx1fl/fl
c
3h after anti-CD3/CD28:
CD69+ cells (%)
CD25+ cells (%)
CD44+ cells (%)
Vehicle
eATP
eATP/ P2RX4i
CD4-Cre
CD4-Cre Panx1fl/fl
d
P14 cells, LCMV-Arm d7 p.i.:
Spleen P14 cells/mouse
sgCd4
sgP2rx4
e
72h after anti-CD3/CD28:
CTV-divided CD8+ T cells (%)
WT
P2RX7-KO
f
WT
Panx1-KO
Effector-like or Memory-like
WT S.N.
KO S.N.
WT I.C.
KO I.C.
GC-MS Metabolomics
g
Effector-like i.c.
Effector-like s.n.
Memory-like i.c.
Memory-like s.n.
Scores Plot
h
Effector-like
Memory-like
row min
row max
Top 100 DEMs
WT > Panx1-KO s.n.
WT_001
WT_002
WT_003
KO_001
KO_002
KO_003
PE(15:0/20:2(11Z,14Z))
Gly Phe
LysoPC(16:0)
Cytosine
L-Lactate
Muramic acid
Pro Pro Pro
Tremetone
Adenine
Ile Trp Asp
PE(O-20:0/17:2(9Z,12Z))
C18:1n-13
i
Effector-like CD8+ T cells
mRNA (counts per million)
Slc16a1
Slc16a3
ns
****
k
% of total PIs
Saturated
Polyunsaturated
CD4-Cre
CD4-Cre Panx1fl/fl
j
DEMs: WT > Panx1-KO s.n.
Rescue of Panx1-KO CD69+ (%)
Vehicle
Hydroxyectonine
Gly-Phe
N-Acetyl L-Tyrosin EEM
NEAAs
4-Aminopyridine
4-Imidazoleacetic acid
5-Methylcytidine
N4-Acetylcytidine
O-AcetylSerine
Cytosine
C3MP
Hydroxylauric Acid
(Sodium) Lactate
Sulfapyridine
Alanine
Arginine
Asparagine
Cysteine
Gly His
Glycine
Guanosine
Levulinic
Proline
Serine
Tyrosine
Valine
l
72h after anti-CD3/CD28:
Cholera Toxin B gMFI
Sodium lactate
MCT1i
CD4-Cre
CD4-Cre Panx1fl/fl
m
Oleic Acid (a.u.)
CD4-Cre
CD4-Cre Panx1fl/fl
n
72h after anti-CD3/CD28:
Cholera Toxin B gMFI
Sodium lactate
CDIPTi

### Figure S6

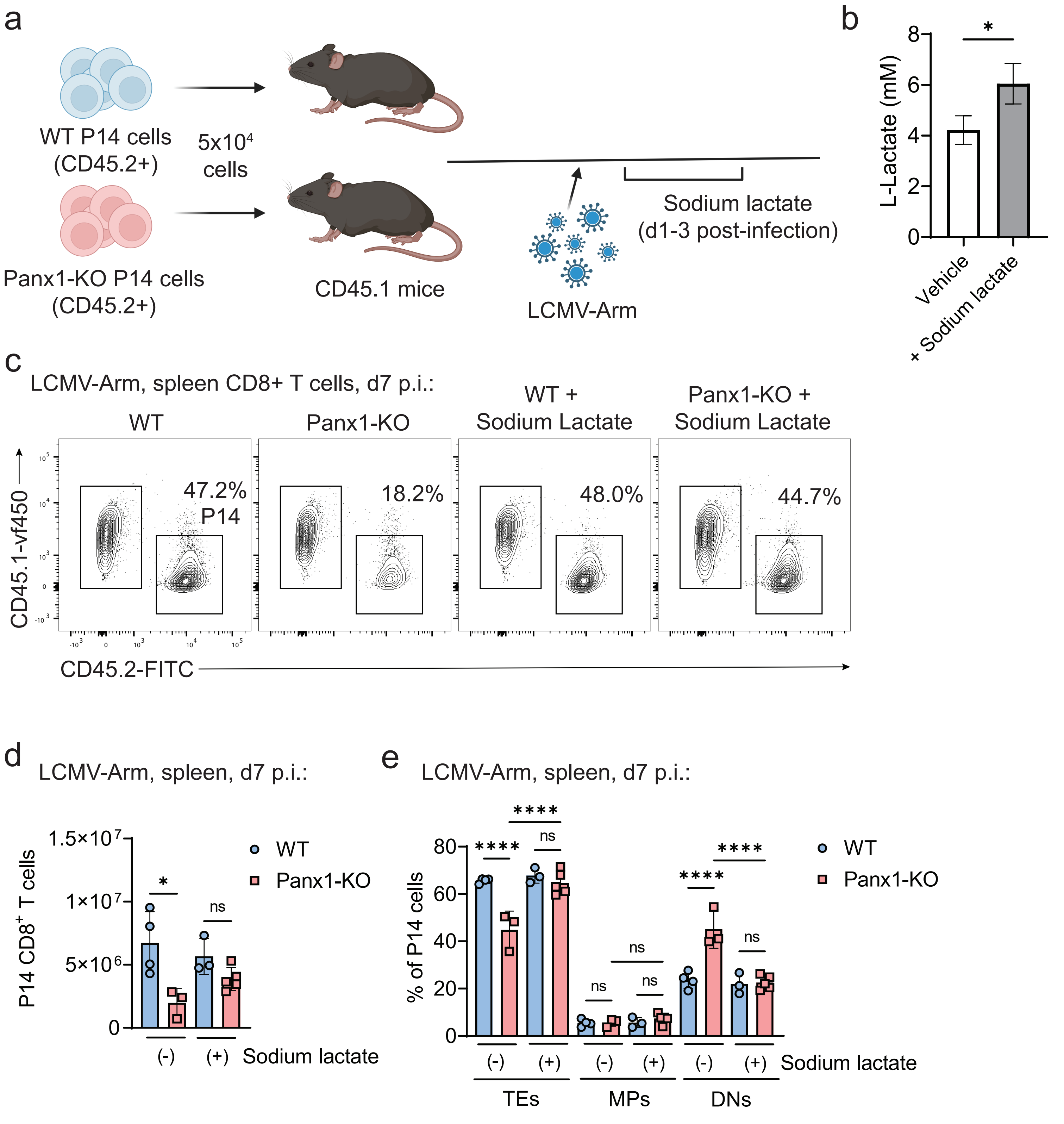

a
WT P14 cells
(CD45.2+)
5x10⁴
cells
Panx1-KO P14 cells
(CD45.2+)
CD45.1 mice
LCMV-Arm
Sodium lactate
(d1-3 post-infection)
b
L-Lactate (mM)
Vehicle
+ Sodium lactate
c
LCMV-Arm, spleen CD8+ T cells, d7 p.i.:
WT
Panx1-KO
WT +
Sodium Lactate
Panx1-KO +
Sodium Lactate
47.2%
P14
18.2%
48.0%
44.7%
CD45.1-vf450
CD45.2-FITC
d
LCMV-Arm, spleen, d7 p.i.:
P14 CD8+ T cells
WT
Panx1-KO
(-)
(+)
Sodium lactate
e
LCMV-Arm, spleen, d7 p.i.:
% of P14 cells
WT
Panx1-KO
(-)
(+)
Sodium lactate
TEs
MPs
DNs

### Figure S7

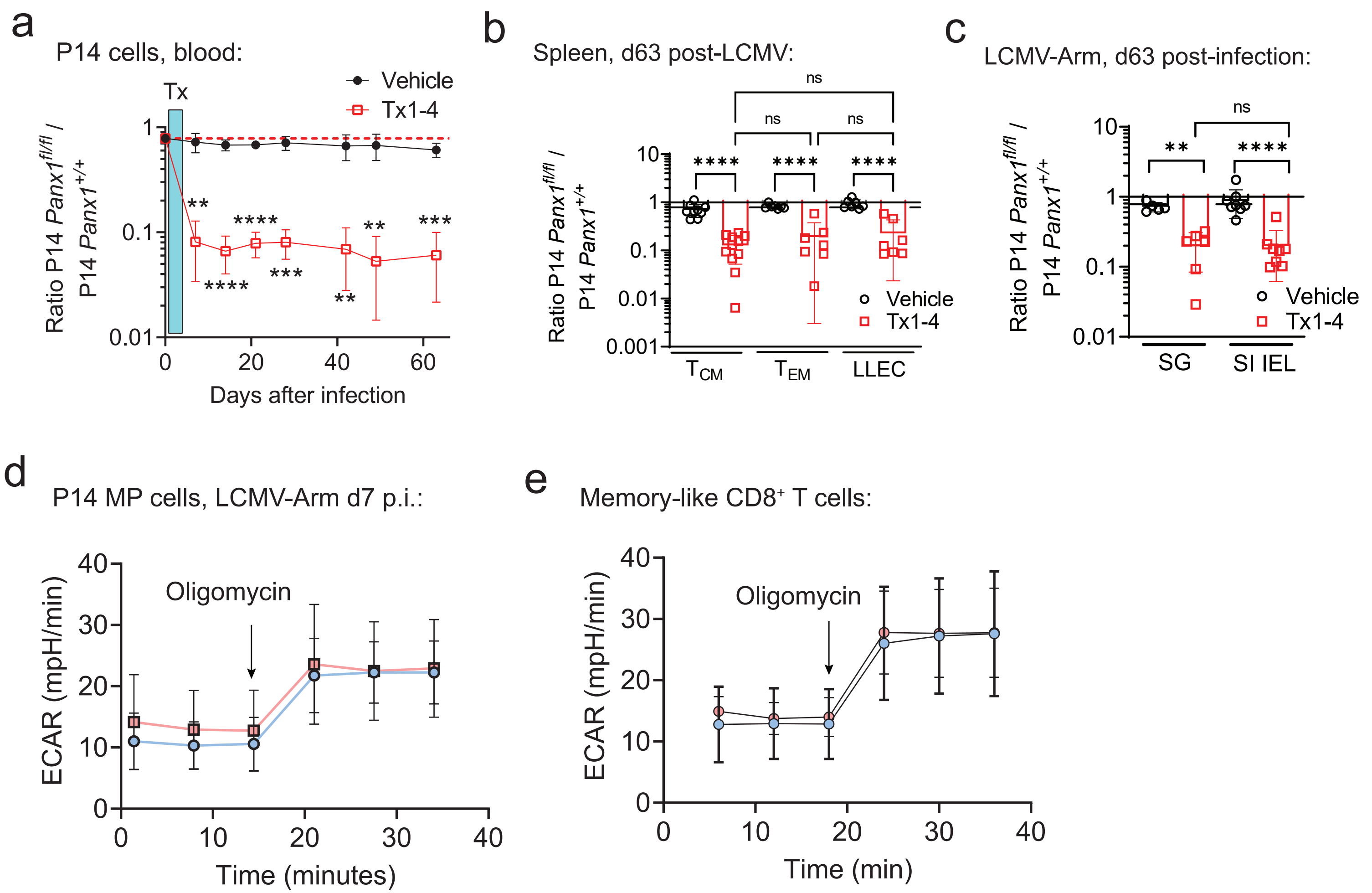
a
P14 cells, blood:
Tx
Vehicle
Tx1-4
Ratio P14 Panx1fl/fl / P14 Panx1+/+
Days after infection
b
Spleen, d63 post-LCMV:
ns
Vehicle
Tx1-4
TCM
TEM
LLEC
c
LCMV-Arm, d63 post-infection:
SG
SI IEL
d
P14 MP cells, LCMV-Arm d7 p.i.:
Oligomycin
ECAR (mpH/min)
Time (minutes)
e
Memory-like CD8+ T cells:
Oligomycin
ECAR (mpH/min)
Time (min)
